# Acetylcholine mediates behavioral state-dependent modulation of spontaneous activity in the developing mouse visual cortex

**DOI:** 10.64898/2026.08.30.748114

**Authors:** David Cabrera-García, Jude Abeje, Christian Lohmann

## Abstract

Synchronized and desynchronized patterns of spontaneous activity during brain development prepare cortical circuits such as the visual cortex for processing external sensory information. Yet the neuromodulatory mechanisms that regulate these early activity patterns remain poorly understood. One such neuromodulator is acetylcholine, which tracks behavioral states and modulates visual cortical activity in adults, but whose function in the developing cortex is largely unexplored. Here, we combined in vivo imaging of genetically encoded acetylcholine and calcium sensors with movement recordings to characterize the relationship between cholinergic dynamics, neuronal activity, and behavioral states in the mouse visual cortex during the second postnatal week, before eye opening. We found that acetylcholine release in the primary visual cortex (V1) closely tracked body movements but was not evoked by visual stimulation alone. Periods of elevated acetylcholine were associated with reduced pairwise neuronal correlations, longer network events with greater temporal jitter, and a transient reduction in mean V1 activity. Chemogenetic inhibition of basal forebrain cholinergic neurons increased the internal synchrony of network events and partially reversed the movement-associated reduction in V1 activity, identifying basal forebrain input as a source of this cholinergic modulation. These findings establish acetylcholine as a state-dependent regulator of spontaneous activity in the developing visual cortex before visual experience begins.

## INTRODUCTION

Spontaneous neuronal activity during brain development is required for circuit maturation and prepares sensory cortical areas such as the visual cortex for sensory processing ^1–4^. In the visual system, much of this activity occurs in utero in humans, while in rodents it continues over the first two postnatal weeks until eye opening. During this period, spontaneous correlated firing in retinal ganglion cells, known as retinal waves, refines the visual pathway in the dorsal lateral geniculate nucleus, superior colliculus, and visual cortex ^5–10^. In parallel, the visual cortex becomes less sensitive to retinal waves ^11^, and cortical activity transitions from highly synchronized to desynchronized ^4^, preparing the circuit for visual processing at eye opening ^12^. However, how spontaneous cortical activity is modulated across this developmental period remains incompletely understood ^4^. In particular, the function of neuromodulators such as acetylcholine, which strongly shapes visual cortical activity across behavioral states in adults ^13^, remains largely unexplored in the developing visual cortex.

Acetylcholine is a neuromodulator of particular interest because its cortical levels fluctuate across the sleep-wake cycle, rising during arousal ^14–16^ and rapid eye movement (REM) sleep ^17,18^. Across the dorsal cortex, acetylcholine and neuronal calcium dynamics are co-modulated by locomotion and arousal, with acetylcholine more tightly coupled to locomotion frontally and calcium activity preferentially coupled to arousal in posterior areas ^14^. In the adult visual cortex, activity of cholinergic axons from the basal forebrain, the main source of acetylcholine in the cortex, also closely tracks whisking ^15,19,20^ and locomotion ^15,21^. In addition, cholinergic input from the basal forebrain mediates decorrelation of V1 neuronal activity during locomotion ^21^, and this decorrelation enhances visual responses ^22,23^. These findings link acetylcholine release in the adult visual cortex to active behavioral states and to fluctuations in neuronal activity.

Although recent work has identified desynchronizing effects of the neuromodulators serotonin and oxytocin in the developing somatosensory and visual cortices ^24,25^, much less is known about the function of acetylcholine in the developing cortex before eye opening. Anatomically, cholinergic axons from the basal forebrain already reach the cortex during the first postnatal days ^26^, and both nicotinic and muscarinic acetylcholine receptors are expressed in the cortex by the second postnatal week in rodents ^27^. Functionally, muscarinic agonists trigger spontaneous activity waves in neocortical slices at postnatal day 5 (P5) ^28^. In vivo, pharmacologically increasing acetylcholine levels in V1 reduces spindle bursts, whereas blocking muscarinic signaling has the opposite effect at P5-6 ^29^. Previous studies in neonates, however, have not linked cholinergic signaling to the behavioral states that modulate cortical activity at this age. Sleep, for instance, predominates in neonates and may be pivotal for spontaneous activity-dependent development ^30^. Movements during wakefulness, in turn, are associated with a reduction in spontaneous cortical activity during the first postnatal weeks, including in the visual cortex ^31,32^. The wake-related reduction in activity is specific to development and reverses after eye opening ^32^, when locomotion desynchronizes neuronal activity ^21,33^ and increases visually evoked firing rates ^34^. These observations point to an unexplored developmental role for acetylcholine in the state-dependent modulation of spontaneous activity. Directly testing this idea has been difficult, however, because real-time acetylcholine and neuronal dynamics are challenging to measure in neonates. The recent development of genetically encoded acetylcholine sensors ^35^ now makes such in vivo experiments possible.

Here, we investigated the function of acetylcholine in the developing visual cortex before eye opening using in vivo two-photon and widefield imaging with genetically encoded sensors for acetylcholine and calcium, while inferring behavioral states from body and facial movements. We show that acetylcholine signals are present during the second postnatal week, increase at the onset of body movements associated with active sleep and wakefulness, and are not driven by visual stimulation alone. Elevated acetylcholine is associated with desynchronized cortical activity and altered temporal network-event structure. Finally, using chemogenetics, we identify basal forebrain cholinergic input as a source of these state-dependent changes in V1 activity.

## RESULTS

### Acetylcholine signaling is present in the visual cortex at the beginning of the second postnatal week

To study acetylcholine and neuronal dynamics in the developing visual cortex before eye opening, we injected the green GRAB (G-protein-coupled receptor-activation-based) acetylcholine sensor GRAB_ACh3.0_ (GACh) ^35^ and the red calcium indicator jRGECO1a ^36^, both under the synapsin promoter, into the visual cortex of P0-1 mice (**Fig. 1A**).

**Figure 1.**
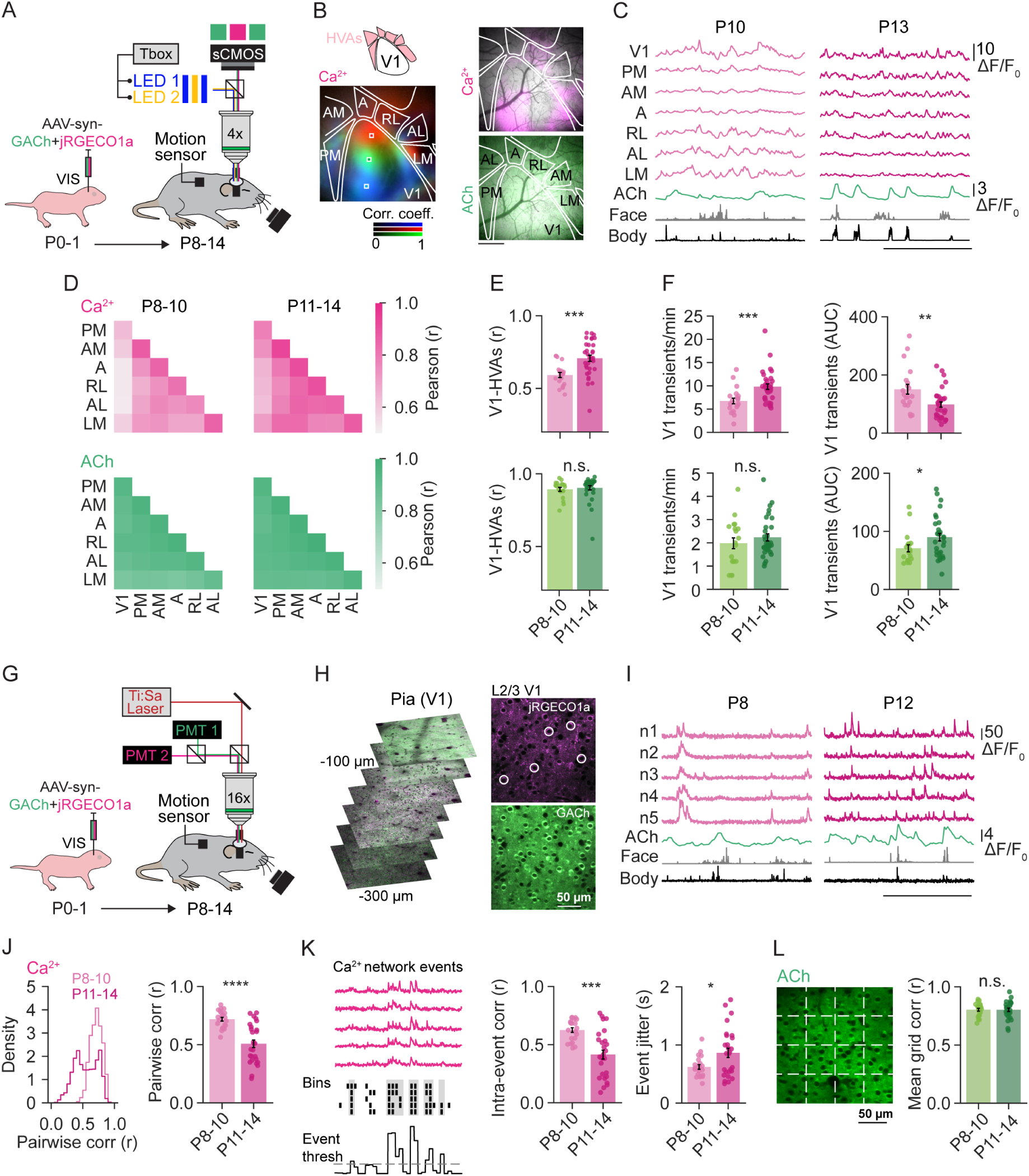
Postnatal acetylcholine signaling increases during the transition to sparse neuronal activity in the mouse visual cortex before eye opening. **A.** Neonatal mice were injected in the visual cortex (VIS) at P0-1 with AAVs encoding a red calcium sensor (jRGECO1a) and a green acetylcholine sensor (GACh) under the synapsin promoter. Widefield imaging was performed in unanesthetized, head-fixed mice at P8-14. A custom trigger box (see Methods) sequentially triggered LEDs to acquire both signals across visual cortical areas while face and body movements were recorded. **B.** Left: Functional correlation maps were generated from correlations between the time courses of selected seed regions in V1 and higher visual areas (HVAs). The final RGB map reflected the topographic organization of spontaneous cortical activity. Right: Example frames of a spontaneous calcium network event (top) and the corresponding acetylcholine signal (bottom). HVA abbreviations: PM (posteromedial), AM (anteromedial), A (anterior), RL (rostrolateral), AL (anterolateral), LM (lateromedial). Scale bar: 0.5 mm. **C.** Representative traces at P10 (left) and P13 (right) showing mean neuronal activity (magenta) from V1 and HVAs, acetylcholine in V1 (green), face camera (gray), and motion sensor (black) signals. Horizontal scale bar: 60 s. **D.** Mean functional correlations (Pearson r) between V1 and HVAs for calcium (top) and acetylcholine (bottom) at P8-10 and P11-14. **E.** The correlation of neuronal activity between V1 and HVAs increased during the second postnatal week. The correlation of acetylcholine signals did not change during this period. **F.** Left: The frequency of calcium transients in V1 increased, whereas the frequency of acetylcholine transients remained unchanged between P8-10 and P11-14. Right: The area under the curve (AUC) of calcium transients in V1 decreased, whereas the AUC of acetylcholine transients increased. **G.** Schematic of the two-photon imaging setup for simultaneous imaging of calcium and acetylcholine sensors, and the recording of face and body movements. **H.** Left: Representative two-photon images spanning from the pia to 300 μm below the cortical surface, showing broad expression of GACh and jRGECO1a in V1 under the synapsin promoter. Right: L2/3 neurons in V1 showing calcium sensor (top) and acetylcholine sensor (bottom) signals from the same field of view. **I.** Calcium traces from a representative subset of L2/3 neurons in a field of view similar to that in H, together with the acetylcholine signal averaged across the entire field of view. Bottom traces show face and body movements recorded in parallel. Horizontal scale bar: 60 s. **J.** Pairwise correlations (pairwise corr) of L2/3 neuronal activity in V1 decreased from P8-10 to P11-14, reflecting the progressive desynchronization of cortical activity. **K.** Left: Schematic of network event detection showing a subset of neurons from a FOV similar to **H**. Network events were defined as epochs with more than 20% neuronal participation within a 2-s sliding window. Detected network events are shaded in gray. Middle: Pairwise correlations within network events decreased between P8-10 and P11-14. Right: Network-event jitter increased between P8-10 and P11-14. **L.** Spatial homogeneity of the acetylcholine signal across the two-photon field of view did not differ between P8-10 and P11-14. Data are shown as mean +/- SEM. Each dot represents the mean for one animal. **D-F**: Acetylcholine (P8-10: 19 mice; P11-14: 31 mice), calcium (P8-10: 20 mice; P11-14: 33 mice); **J-K** (P8-10: 21 mice; P11-14: 27 mice); **L** (P8-10: 21 mice; P11-14: 26 mice). Statistical comparisons used unpaired t-tests or Mann-Whitney tests. n.s., not significant, \**P* < 0.05, \*\**P* < 0.01, \*\*\**P* < 0.001, \*\*\*\**P* < 0.0001.

We first verified the specificity of the GACh signal during the second postnatal week. In acute brain slices, confocal imaging revealed a strong increase in fluorescence upon application of acetylcholine and muscarine, whereas glutamate did not evoke a response (**Fig. S1A-B**). Consistent with GACh being derived from a muscarinic receptor ^35^, atropine blocked the response to acetylcholine (**Fig. S1C**). In vivo, 1 mM acetylcholine puffed onto superficial V1 evoked a similarly strong response, while the insensitive GACh-mut sensor did not (**Fig. S1D**). Thus, GACh was robustly expressed and responded specifically to cholinergic stimulation during the second postnatal week.

We then performed acute in vivo widefield (**Fig. 1A**) and two-photon imaging (**Fig. 1G**) between P8 and P14 to simultaneously monitor acetylcholine signaling (ACh) and neuronal activity (Ca^2+^) in unanesthetized neonatal mice at the network and cellular levels across behavioral states. For widefield imaging, a custom Arduino trigger box sequentially triggered two excitation LEDs, and a single camera collected both emission signals through a dual bandpass filter. A motion sensor and infrared camera simultaneously recorded face and body movements to infer behavioral states (**Fig. 1A**, see Methods). With this approach, we acquired acetylcholine and calcium signals from V1 and higher visual areas (HVAs) with no observable crosstalk between green and red channels (**Fig. 1B**). Visual maps from adult mice ^37^ were adjusted using functional correlation maps of spontaneous activity ^38^ (**Fig. 1B**) to extract mean neuronal activity from each visual area (**Fig. 1C**).

To investigate acetylcholine and neuronal dynamics across the second postnatal week, we compared two age groups (P8-10 and P11-14) based on the transition around P11 toward greater desynchronization within V1 and stronger functional correlations among visual areas (**Fig. S1E, G**), consistent with previous studies ^38,39^. Whereas V1-HVA functional correlations increased during the second postnatal week, as recently reported ^38^, acetylcholine signals were highly homogeneous and synchronized across visual areas from the beginning of this period (**Fig. 1D-E**). The frequency of calcium transients in the mean V1 activity (V1 calcium transients) increased, while that of acetylcholine transients remained relatively stable over the same period (**Fig. 1F**). In contrast, the area under the curve (AUC) of V1 calcium transients decreased, whereas the AUC of acetylcholine transients increased slightly (**Fig. 1F**).

To simultaneously measure acetylcholine and neuronal activity at cellular resolution in V1, we recorded from neurons in layers 2 and 3 (L2/3) using two-photon time-lapse microscopy (**Fig. 1G-H**). The synapsin promoter drove broad expression of GACh and jRGECO1a in V1 L2/3 neurons (**Fig. 1H**). Between P8-10 and P11-14, we observed the expected shift from synchronized to decorrelated activity in L2/3 neurons (**Fig. 1I-J, Fig. S1G**) ^12,40^. Synchronous network events, defined by more than 20% neuronal participation, showed weaker intra-event pairwise correlations and greater temporal jitter but similar duration toward the end of the second postnatal week (**Fig. 1K, Fig. S1H**). To test whether the acetylcholine signal was also spatially homogeneous under two-photon imaging, we divided each field of view (FOV) into a grid of ROIs and calculated pairwise correlations between their mean GACh traces ^41^. Correlations were high across ROIs (**Fig. 1L, Fig. S1I-J**), compatible with volume release or synchronized synaptic release in V1 before eye opening, although the spatial resolution of the sensor and imaging setup cannot rule out a local synaptic contribution ^42^.

Together, these results show that the acetylcholine signal is spatially homogeneous across the visual cortex and modestly increases in V1 as neuronal activity becomes sparser and functional correlations between visual areas mature.

### Acetylcholine release in the developing visual cortex is related to movements during active sleep and wakefulness

In adults, acetylcholine is a widespread cortical neuromodulatory signal that tracks whisking and locomotion in V1 ^14,15,19,21^. To test whether cholinergic release is coupled to different movements before eye opening, we aligned the widefield acetylcholine signal in V1 to the onset of brief body movements (twitches) and prolonged movements (wake movements). To isolate movement-associated acetylcholine signals, we analyzed only body twitches and wake movements preceded by a 2-s period of no movement (**Fig. 2A**). Increases in the acetylcholine signal followed both twitches and wake movements at P8-10 and P11-14, and were larger and more sustained during wake movements (**Fig. 2B**). Consistent with the overall increase in the acetylcholine signal across the second postnatal week (**Fig. 1F**), movement-associated responses were also larger at P11-14 (**Fig. 2C**). These responses were not caused by motion artifacts, as the mutant sensor (GACh-mut) showed no increase at movement onset (**Fig. 2D**).

**Figure 2.**
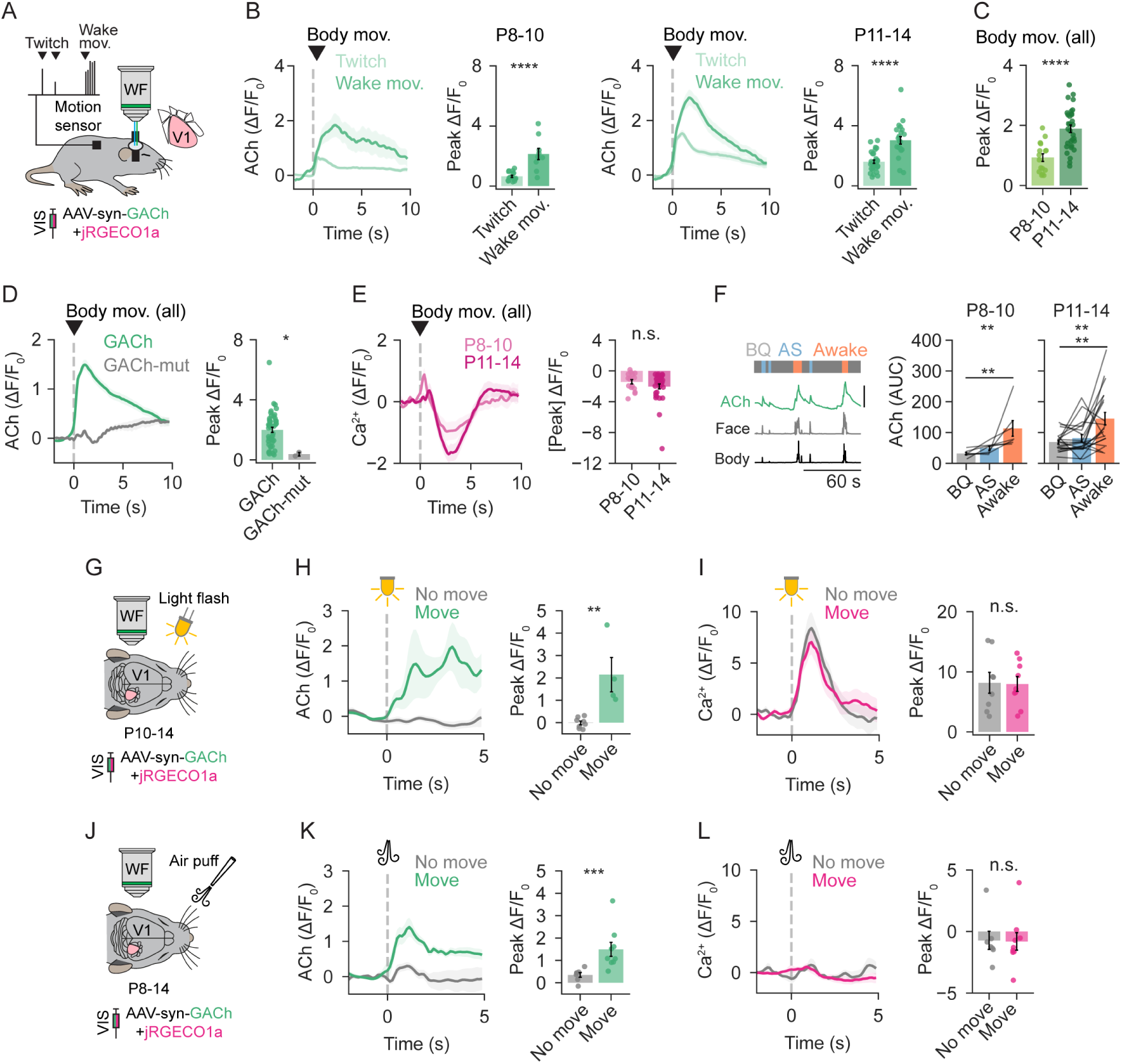
Acetylcholine in the developing V1 tracks spontaneous body movements but not visual stimulation. **A.** Schematic of widefield imaging of the acetylcholine sensor GACh in V1 with simultaneous motion-sensor recording of body movements. Movements were classified as twitches and wake movements. **B.** Peri-event time histogram aligned to the onset of body twitches and wake movements at P8-10 and P11-14, and bar plots with mean peak amplitudes per animal. Wake movements (P8-10: 9 mice; P11-14: 22 mice) elicited larger responses than twitches (P8-10: 16 mice; P11-14: 31 mice). **C.** Acetylcholine signal associated with movements (twitches and wake movements pooled) increased during the second postnatal week (P8-10: 16 mice, P11-14: 31 mice). **D.** No increase in fluorescence was detected at the onset of body movements with GACh-mut (2 mice). **E.** Peri-event time histogram of mean V1 widefield calcium activity aligned to the onset of body movements (twitches and wake movements pooled) at P8-10 (17 mice) and P11-14 (33 mice). **F.** Behavioral states (behavioral quiescence, BQ; active sleep, AS; wakefulness) were defined from face and body movement recordings (see Methods). Acetylcholine levels in V1 were higher during wakefulness than during AS or BQ at both P8-10 (7 mice) and P11-14 (20 mice). Vertical scale bar: 5 ΔF/F_0_. **G.** Contralateral eye stimulation with light flashes in P10-14 mice. **H.** The acetylcholine signal increased after light flashes only when movements followed the stimulus (no movement: 8 mice, movement: 4 mice). **I.** Light flashes evoked robust calcium responses in V1 regardless of whether movements followed the stimulus (no movement: 8 mice; movement: 9 mice). **J.** Contralateral whisker pad stimulation with air puffs in P8-14 mice. **K.** Whisker stimulation evoked only a small acetylcholine increase in V1 when the air puff did not trigger body movements (no movement: 7 mice; movement: 9 mice). **L.** Whisker stimulation did not elicit detectable neuronal activity in V1, regardless of whether body movements occurred (no movement: 7 mice; movement: 9 mice). Data are shown as mean +/- SEM. Signals of peri-event and peri-stimulus time histograms were normalized to a 2-s pre-event baseline, and corresponding peak amplitudes were measured within the first 5 s after movement or stimulus onset. Each dot represents the mean from a single animal. Statistical comparisons between two groups were made using unpaired t-tests or Mann-Whitney U tests, except in **F** (Friedman test with Dunn post hoc comparisons). n.s., not significant; \**P* < 0.05, \*\**P* < 0.01, \*\*\**P* < 0.001, \*\*\*\**P* < 0.0001.

Because the initial acetylcholine kinetics were comparable after twitches and wake movements, we pooled the two movement types to examine the corresponding movement-locked neuronal responses. In contrast to the acetylcholine signal, V1 neuronal activity decreased relative to the 2-s pre-movement baseline following movement onset (**Fig. 2E**), an effect previously reported in V1 that reverses at eye opening ^32,43^. Full-face movements occurring independently of body movements were also associated with increases in acetylcholine release and decreases in V1 activity (**Fig. S2A-C**). The absence of a clear difference between facial twitches and prolonged facial movements may be explained by the limited temporal resolution of the face camera for resolving brief twitches. Although the signal-to-noise ratio was lower, the movement-associated increase in acetylcholine and decrease in calcium activity were also observed by two-photon imaging after pooling body movements (**Fig. S2D-F**).

We then used body twitches and wake movements to infer active sleep (AS) and wakefulness (awake), respectively (see Methods), as these movements provide a proxy for defining behavioral states in pups ^30,44^. Remaining quiet periods were classified as behavioral quiescence (BQ), because quiet sleep and quiet wakefulness cannot be reliably distinguished before P11 without electrophysiological recordings ^44^. Acetylcholine levels differed across behavioral states (**Fig. 2F**), with higher levels during wakefulness throughout the second postnatal week. Acetylcholine levels during AS periods were similar to those during BQ (**Fig. 2F**), likely because AS includes inter-twitch intervals that capture broader temporal dynamics than the immediate response to twitch onset.

A few days before eye opening, intense light stimulation can evoke early visual responses in V1 ^12^, and visual stimulation can evoke acetylcholine release in adult mice ^45,46^, prompting us to examine cholinergic and neuronal responses to whole-field light flashes (100 ms) before eye opening (**Fig. 2G**). Light flashes did not evoke acetylcholine release in V1 on trials without movement (**Fig. 2H**), even though visually evoked neuronal activity was reliably recorded at P10-14 (**Fig. 2I**). When movements were triggered or occurred spontaneously alongside visual stimulation, acetylcholine release increased, whereas calcium responses were unaffected (**Fig. 2H, I**). We next examined whisker stimulation, given the potential for cross-modal processing between visual and somatosensory areas ^47^ and the fact that acetylcholine release tracks whisking in adult mice ^16,19^ (**Fig. 2J**). Gentle air-puff stimulation of the whiskers evoked only a small increase in acetylcholine release in V1, smaller than that observed when the air puff triggered body movements (**Fig. 2K**). No clear calcium response was observed in V1 regardless of whether movements followed the air puff (**Fig. 2L**), although we confirmed that the same stimulation evoked acetylcholine and calcium responses in the rostrolateral (RL) HVA (**Fig. S2G**), which receives whisker-related sensory information during development ^48^.

These results indicate that acetylcholine closely covaried with body and facial movements in the developing V1, as in adult animals. In contrast to adults, V1 neuronal activity in neonates was transiently reduced following movement onset, and acetylcholine was not evoked by visual stimulation alone.

### Spontaneous activity in V1 is decorrelated during periods of high acetylcholine

Given the well-established role of acetylcholine in desynchronizing cortical activity in the adult brain ^22,23^, we next examined whether acetylcholine has a similar function during development and may thus contribute to the shift toward desynchronization during the days before eye opening ^4^.

In two-photon recordings of acetylcholine and calcium signals from L2/3 neurons, we segmented recordings into periods of baseline and acetylcholine transients (**Fig. 3A**). Comparing the same neurons across both periods revealed a clear decrease in pairwise correlations during acetylcholine transients at both P8-10 and P11-14 (**Fig. 3B**). To determine whether decorrelation scaled with the duration or magnitude of acetylcholine release, we measured pairwise correlations during each acetylcholine transient (**Fig. S3A-B**). Interestingly, the association between acetylcholine and decorrelation was less pronounced for longer and larger acetylcholine signals, and this pattern was similar in both age groups (**Fig. 3C**, **S3A-B**). Based on these results and the time course of acetylcholine release shown in **Fig. 2B**, we divided acetylcholine transient periods into an early phase (the first 5 s) and a delayed phase (the remainder of the transient). Consistent with this time-dependent effect, we observed that network events, as defined in **Fig. 1K**, occurring during the early phase of acetylcholine release lasted longer and showed more temporal jitter than those during baseline (**Fig. 3D**), and these effects were consistent across the second postnatal week (**Fig. S3C**). Intra-event pairwise correlations were similar between events starting during baseline and those starting during acetylcholine transients (**Fig. 3D, S3C**), possibly because the longer network events extend beyond the early phase when acetylcholine-associated decorrelation is strongest.

**Figure 3.**
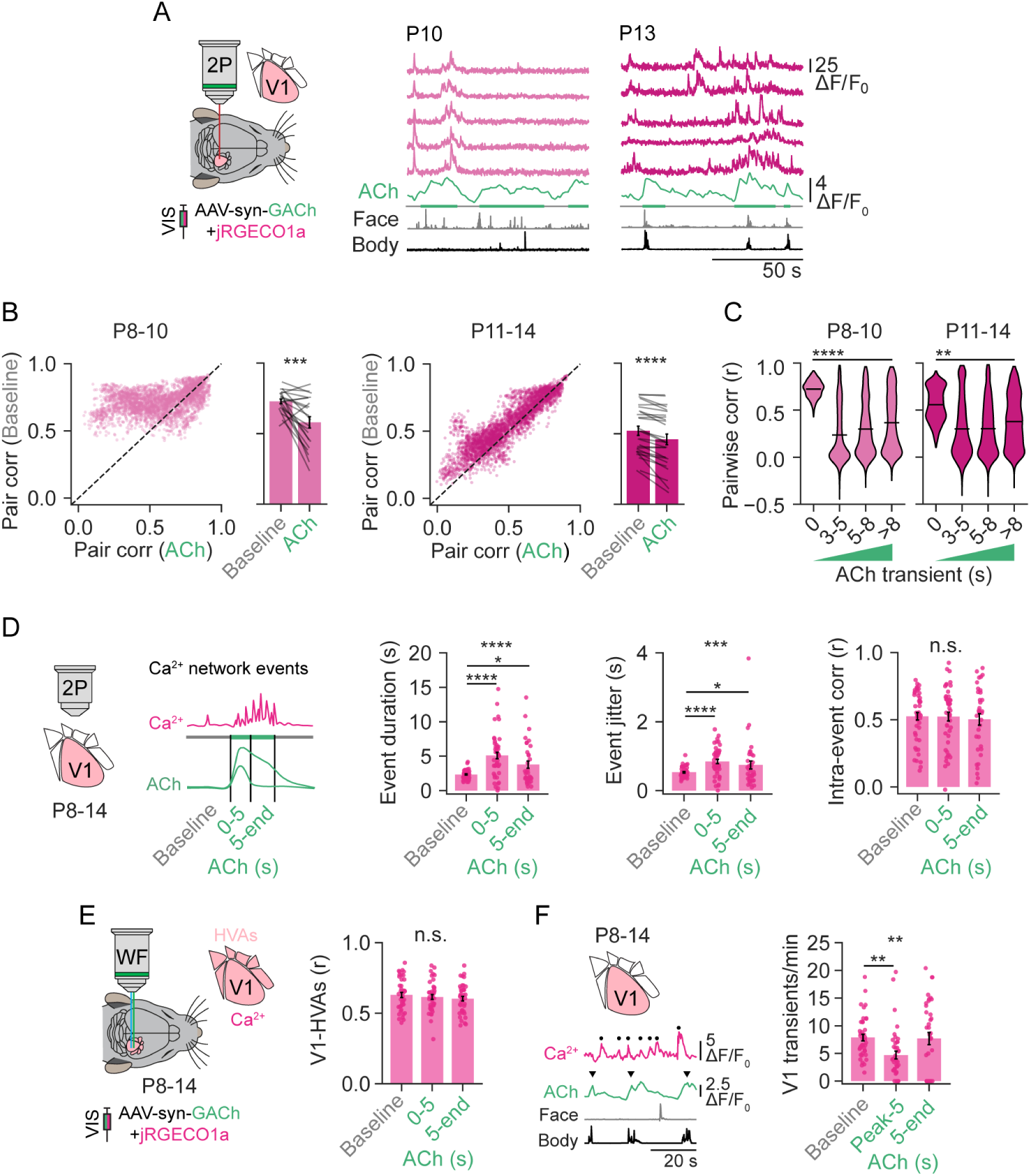
Neuronal activity in V1 is decorrelated during periods of elevated acetylcholine levels. **A.** Simultaneous two-photon imaging of acetylcholine (GACh) and neuronal calcium activity (jRGECO1a) in V1 L2/3 neurons in P8-14 mice. Baseline (horizontal gray bars) and elevated acetylcholine periods (horizontal green bars) were defined based on acetylcholine transients. Right: Example traces from P10 and P13 mice. **B.** Pairwise correlations of neuronal activity in V1 were lower during acetylcholine transients at both P8-10 and P11-14. Each dot in the scatter plot represents the mean pairwise correlation per neuron, and lines in the bar plot represent the mean per animal. **C.** Acetylcholine-associated decorrelation was more pronounced during the first seconds of the acetylcholine transient. Pairwise correlations within each acetylcholine transient were quantified and binned by acetylcholine transient duration (Tukey post hoc comparisons: baseline vs each acetylcholine transient bin). **D.** Schematic (left): Acetylcholine periods were segmented into the first 5 s and the rest of the transient. Bar plots (right): Network events that began during acetylcholine transients had longer durations (left), higher temporal jitter (middle), and similar intra-event pairwise correlations (right) compared to events beginning during baseline periods (post hoc Dunn: baseline vs 0-5; baseline vs 5-end). **E.** Left: Sequential widefield imaging of calcium and acetylcholine signals in mice between P8 and P14. Right: Functional correlations between mean V1 and HVA activity were similar during baseline and elevated acetylcholine periods. HVA activity represents the mean of RL, A, AL, AM, PM, and LM. **F.** Left: Example traces from a P10 mouse depicting V1 calcium (magenta) and acetylcholine (green) signals, together with face (gray) and body (black) movement signals. Vertical scale bars: 5 ΔF/F_0_ (calcium), 2.5 ΔF/F_0_ (acetylcholine). Black dots indicate calcium transients. Black arrowheads indicate acetylcholine peaks. Right: The frequency of V1 calcium transients decreased during the first 5 s after the acetylcholine peak (post hoc Dunn test). Data are shown as mean +/- SEM. Each dot or line represents the mean from a single animal unless stated otherwise. **B-C** (P8-10: 21 mice; P11-14: 27 mice), **D** (baseline: 47 mice; 0-5: 46 mice; 5-end: 35 mice), **E** (baseline: 41 mice; 0-5: 41 mice; 5-end: 37 mice), **F** (baseline: 41 mice; Peak-5: 41 mice; 5-end: 37 mice). Statistical comparisons were made using paired t-tests (**B**) and Kruskal-Wallis or one-way ANOVA with Dunn or Tukey post hoc test, respectively (**C-F**). n.s., not significant; \**P* < 0.05, \*\**P* < 0.01, \*\*\**P* < 0.001, \*\*\*\**P* < 0.0001.

We then used widefield imaging (**Fig. 3E**) to evaluate whether periods of elevated acetylcholine might contribute to establishing proto-sensory maps before the onset of active sensory experience ^49^ by modulating the correlations between visual areas. However, correlations between mean spontaneous activity in V1 and HVAs were similar during baseline and elevated-acetylcholine periods (**Fig. 3E, S3D**). This result may reflect the dominant contribution of thalamic activity to correlations between visual areas at this developmental stage ^38^. Our data (**Fig. 2E**) also confirmed the previously described movement-associated reduction in spontaneous V1 activity ^12,31,50^. We therefore compared the frequency of V1 calcium transients between baseline and the first 5 s after the acetylcholine peak. Consistent with the above results, we found a decrease in V1 transient frequency during the first 5 s after the acetylcholine peak (**Fig. 3F, S3E**).

The time-dependent acetylcholine effects on V1 activity raised the possibility that they reflect the biphasic voltage responses to acute acetylcholine in pyramidal neurons described in adult animals ^51,52^. To test this, we performed patch-clamp recordings from V1 pyramidal neurons in slices. In response to local acetylcholine (1 mM) puffs, approximately half of the neurons showed biphasic responses, with an initial hyperpolarization lasting up to 5 s followed by delayed depolarization (**Fig. S3F, G**). We then applied acetylcholine focally in vivo while imaging neuronal activity with a red calcium indicator using two-photon microscopy and found that activity increased asynchronously in a substantial proportion of neurons after a delay of at least 3 s (**Fig. S3H**). Although 1 mM is higher than estimated extrasynaptic acetylcholine concentrations in the cortex ^42^, these results are consistent with a transient net reduction in V1 activity during the first few seconds after acetylcholine elevation.

The close coupling between acetylcholine transients and active behavioral states prompted us to compare spontaneous V1 activity across states. We found a decrease in the frequency of V1 calcium transients during wakefulness (wake movements) and an increase during AS (**Fig. S3I**). This increase in V1 activity during AS likely reflects neuronal dynamics occurring during the longer intervals between body twitches, which can include bouts of facial twitches. Both active behavioral states were associated with neuronal decorrelation, but only at the end of the second postnatal week (**Fig. S3J**), when movement-associated acetylcholine responses were also larger.

Together, these results show a strong local decorrelation of neuronal activity and a transient reduction in V1 activity during acetylcholine transients, effects that parallel those observed during wakefulness. The biphasic voltage responses of pyramidal neurons to acetylcholine may partly account for the transient reduction in V1 activity.

### Cholinergic signaling from the basal forebrain mediates modulation of spontaneous activity

Our results so far revealed a strong association of elevated acetylcholine with reduced neuronal correlations, altered network event structure (**Fig. 3A-D**), and a transient decrease in mean activity in V1 (**Fig. 3F**). We therefore perturbed endogenous cholinergic signaling pharmacologically and chemogenetically to determine whether it is required for the movement- and state-associated changes in V1 activity and network structure. Given the similar acetylcholine-associated effects on neuronal activity at P8-10 and P11-14 (**Fig. S3A-E**), we did not test age-dependent differences in the following experiments.

First, we focally applied atropine (1 mM) to the surface of the visual cortex to block muscarinic receptors, thereby reducing the acetylcholine sensor signal in V1 (**Fig. 4A, B**). This was associated with an increase in the frequency of V1 calcium transients (**Fig. 4A**). At the network level, atropine disrupted the functional retinotopy within V1 and between visual areas by increasing correlations between V1 and HVA activity (**Fig. S4A-B**), likely due to the marked increase in synchronous activity within V1. Locally, atropine application strongly increased both full-trace and intra-event pairwise correlations between neurons in L2/3 of V1 (**Fig. 4B-C, S4C**), including during pooled AS and wakefulness periods (**Fig. S4D**). The duration and jitter of network events also decreased (**Fig. 4C, S4E**). These results support a requirement for cortical muscarinic signaling in desynchronizing spontaneous network activity during the second postnatal week.

**Figure 4.**
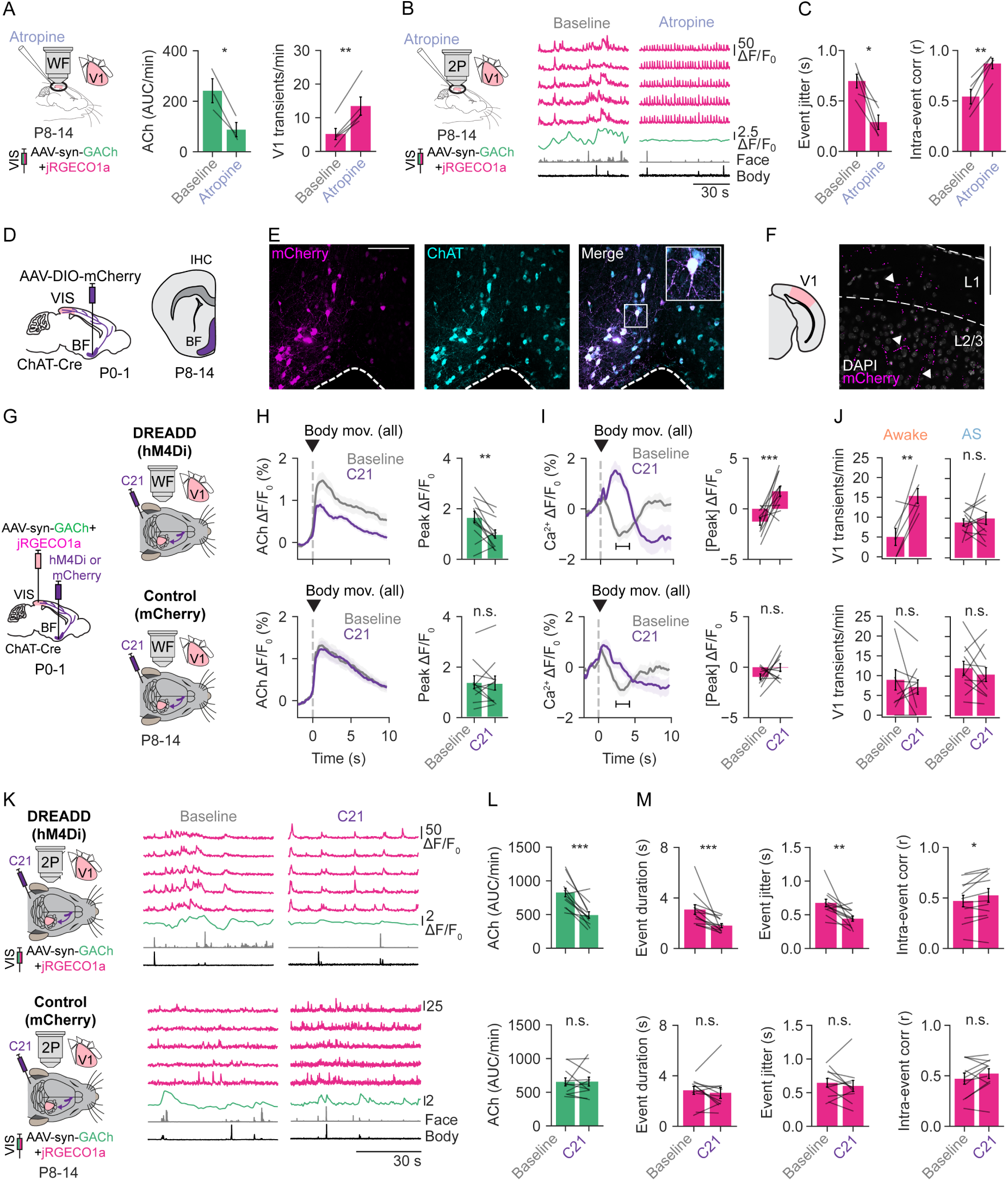
Basal forebrain cholinergic signaling modulates spontaneous activity in the developing mouse visual cortex. **A.** Left: Local application of atropine reduced the signal of the acetylcholine sensor during widefield imaging (3 mice). Right: The frequency of V1 calcium transients increased after application of atropine (5 mice). **B.** Example traces from L2/3 neurons in V1 during baseline and after local cortical application of atropine (P13 mouse). **C.** Focal application of atropine in V1 decreased event jitter (left) and increased intra-event pairwise correlations (right). **D.** ChAT-Cre mice were injected with AAV-DIO-mCherry into the basal forebrain, and immunohistochemistry was performed between P8 and P14. The basal forebrain (BF) is highlighted in magenta. **E.** Representative confocal images from a P13 mouse. mCherry expression colocalized with ChAT-positive neurons, confirming cholinergic specificity. Scale bar: 200 μm. **F.** Injection of AAV-DIO-mCherry into the basal forebrain of ChAT-Cre mice at P0-1 labeled cholinergic axonal projections within V1. Example confocal image from a P13 mouse. White arrowheads indicate labeled axonal varicosities. Scale bar: 100 μm. **G.** Experimental design for Cre-dependent expression of hM4Di or mCherry in the basal forebrain of ChAT-Cre mice (P0-1), combined with pan-neuronal expression of GACh and jRGECO1a in the visual cortex. Widefield imaging was performed before and after subcutaneous injection of the DREADD agonist C21 in hM4Di and mCherry control mice. **H.** Movement-associated acetylcholine signals were reduced following C21 administration in hM4Di mice but not in mCherry controls. **I.** The movement-onset reduction in calcium activity was partially reversed by chemogenetic reduction of acetylcholine levels in hM4Di mice, but not in mCherry controls. The horizontal bar indicates the window (2-4 s) within which the amplitude was measured. **J.** Frequency of V1 calcium transients during wakefulness (left panels) and AS (right panels) before and after C21 injection in hM4Di (awake: 6 mice, AS: 12 mice) and mCherry mice (awake: 10 mice, AS: 11 mice). **K.** Representative simultaneous two-photon GACh and jRGECO1a recordings from hM4Di and mCherry mice before and after C21 injection. **L.** Acetylcholine levels were reduced in hM4Di mice but not in control mice after C21 injection. **M.** Chemogenetic inhibition of basal forebrain cholinergic neurons resulted in shorter network events with lower temporal jitter and higher intra-event pairwise correlations, indicating increased internal synchrony. Data are shown as mean +/- SEM. Each dot or line represents the mean from a single animal. **C** (5 mice), **H**-**I, L-M**: hM4Di (12 mice; 5 at P8-10, 7 at P11-14), and mCherry (11 mice; 5 at P8-10, 6 at P11-14). Statistical comparisons before and after C21 administration within each group were made using paired t-tests or Wilcoxon signed-rank tests. n.s., not significant; \**P* < 0.05, \*\**P* < 0.01, \*\*\**P* < 0.001, \*\*\*\**P* < 0.0001.

To reduce endogenous acetylcholine without directly blocking cortical receptors and to specifically target the basal forebrain, the primary source of acetylcholine in the visual cortex ^53,54^, we virally expressed a Cre-dependent inhibitory DREADD in choline acetyltransferase (ChAT)-Cre mice.

First, we validated the viral transduction approach in neonates by injecting Cre-dependent fluorescent proteins into the basal forebrain of ChAT-Cre mice and performing immunohistochemistry. Injections at P0-1 successfully targeted the basal forebrain (**Fig. 4D-E**, **S4F**), and over 90% of fluorescent protein-positive neurons in the basal forebrain were also ChAT-positive (**Fig. S4G**). Cholinergic axons arising from basal forebrain neurons were already present in V1 during the second postnatal week (**Fig. 4F, S4H**) and were co-labeled by the ChAT antibody (**Fig. S4I**). Furthermore, ChAT-Cre-dependent expression and ChAT immunohistochemistry showed that the density of cholinergic neurons in the basal forebrain and of cholinergic axons in V1 did not significantly change over the second postnatal week (**Fig. S4J-K**).

Next, we expressed acetylcholine and calcium sensors in the visual cortex at P0-1 and a Cre-dependent inhibitory DREADD (hM4Di) or a fluorescent protein (mCherry as control) in the basal forebrain (**Fig. 4G**). In vitro, bath application of the DREADD agonist C21 ^55^ reduced evoked firing in hM4Di-expressing cholinergic neurons in basal forebrain slices (**Fig. S4L-M**). In vivo, the efficacy of the inhibitory DREADD was confirmed by a reduction in movement-associated acetylcholine responses (**Fig. 4H**), and by an overall 50% reduction in the acetylcholine signal in hM4Di mice, but not in mCherry controls, after C21 injection (**Fig. S4N**).

Chemogenetically reducing acetylcholine release in the cortex partially reversed the movement-associated reduction in neuronal activity (**Fig. 4I**), without affecting the occurrence of movements (**Fig. S4O**). Because mean V1 activity was modulated during acetylcholine transients and active behavioral states (**Fig. 3F** and **S3I**), we compared the frequency of V1 calcium transients before and after C21 injection during wakefulness and AS. Chemogenetic inhibition increased the frequency of V1 calcium transients during wakefulness but produced no change during AS (**Fig. 4J**), consistent with the wake-specific reduction of neuronal activity seen in **Fig. S3I**. These results suggest that basal forebrain cholinergic input partially mediates the movement-associated reduction in neuronal activity.

C21 injection also reduced acetylcholine levels in two-photon recordings from mice expressing hM4Di but not in mCherry controls (**Fig. 4K-L**). At the cellular level, we found that activation of the inhibitory DREADD in basal forebrain cholinergic neurons increased intra-event pairwise correlations, while reducing V1 network event duration and jitter (**Fig. 4M**).

Intra-event pairwise correlations were also increased during active sleep and wakefulness in hM4Di mice (**Fig. S4P**). These network effects were observed despite no change in full-trace pairwise correlations after C21 administration (**Fig. S4Q**).

These results show that acetylcholine shapes correlated activity in the developing visual cortex, and that reducing basal forebrain cholinergic input increases the internal synchrony of network events and modulates V1 activity associated with movements.

## DISCUSSION

Here we show that acetylcholine already displays behavioral state-dependent dynamics in the developing mouse visual cortex during the week before eye opening and modulates neuronal correlations and activity in V1. Chemogenetic inhibition of basal forebrain cholinergic neurons partially reversed the changes in V1 activity and network-event structure, identifying the basal forebrain as a source of state-dependent cholinergic modulation at this early stage. We hypothesize that acetylcholine acts on two timescales: transient decorrelation may refine visual circuits and enable activity-dependent plasticity on short timescales, whereas cholinergic modulation across the second postnatal week may contribute to the shift from synchronized to desynchronized cortical activity that prepares V1 for sensory processing at eye opening.

Previous studies provided indirect evidence for cholinergic modulation of cortical activity during the first postnatal week. Muscarinic agonists trigger cortical waves in neonatal slices ^28,56^, and pharmacological manipulation of cholinergic signaling bidirectionally modulates spontaneous spindle bursts in the neonatal rat visual cortex in vivo ^29^. Behavioral state has similarly been proposed as a regulator of early cortical activity, since spontaneous cortical calcium waves are reduced during self-generated movements ^31^, and spike rates and spindle bursts driven by retinal waves are reduced by movement before eye opening ^32,50^.

Taken together, these observations pointed to a link between cholinergic signaling, behavioral state, and early cortical activity, but the in vivo dynamics of acetylcholine were unknown. Our results integrate these lines of work by directly monitoring acetylcholine in unanesthetized mice, showing that cholinergic signaling tracks movements associated with active behavioral states at least a week before eye opening. Consistent with this state dependence, acetylcholine release was higher during prolonged body movements associated with wakefulness than during twitches associated with active sleep. The higher acetylcholine levels during wakefulness likely reflect the prolonged nature of wake movements, as in adult mice ^19^, since the initial rise in acetylcholine was similar for twitches and prolonged body movements. Across the second postnatal week, the increase in movement-related acetylcholine release may reflect a slight expansion of cholinergic innervation reported elsewhere ^26,57^, or be partly attributable to stronger sensor expression at later ages. The movement-associated release parallels the acetylcholine dynamics linked to arousal and spontaneous behaviors in adult mice ^16,58^, suggesting a mechanism that precedes the visually driven acetylcholine responses of the mature visual cortex ^45,46^. Fully disentangling movement from state as drivers of acetylcholine release will require simultaneous recording and manipulation of basal forebrain cholinergic neurons and cortical acetylcholine release across movements and behavioral states.

In association with movement onset and wakefulness, we also observed a transient decrease in V1 activity, consistent with previous work ^31,50^. Endogenous acetylcholine transients were associated with a comparable decrease in V1 activity, and chemogenetic reduction of cortical acetylcholine levels reversed the reduction that followed movement onset, indicating that the wake-related reduction of V1 activity is partially mediated by acetylcholine. In contrast, chemogenetic reduction of cholinergic input did not detectably alter V1 activity during AS, although the twitch-based inference of AS limits interpretation of this negative result. Noradrenergic input from the locus coeruleus is also tightly coupled to movement onset and may contribute to the modulation that remains after chemogenetic inhibition of cholinergic neurons ^16,19^. The functional role of this movement-associated reduction in activity in the developing visual cortex remains uncertain ^32,50^. Acetylcholine might suppress spontaneous cortical activity during motor-related events before maturation of thalamic drive reverses the sign of this modulation ^32^. Previous reports on the influence of twitches on visual cortex activity have been mixed ^50,59^, but the developing visual cortex likely differs from sensorimotor systems, where such twitches drive spontaneous activity ^30,60^.

Beyond this state-dependent modulation of activity levels, our data also address how acetylcholine shapes the correlation structure of V1. Local muscarinic blockade with atropine strongly increased pairwise correlations and disrupted functional retinotopy, and chemogenetic reduction of basal forebrain cholinergic output shortened network events and increased their synchrony even though movement occurrence was preserved. The stronger effect of atropine relative to chemogenetic silencing can be attributed to two factors. First, atropine blocks all subtypes of muscarinic receptors in V1 with high affinity, whereas hM4Di activation only partially reduces acetylcholine release from basal forebrain terminals. Second, the basal forebrain contains functionally distinct subpopulations and circuit motifs ^61^, and partial silencing may therefore alter neuronal ensembles by disrupting the spatiotemporal structure of endogenous cortical acetylcholine release ^42^, a possibility that our imaging resolution cannot resolve. Consistent with this, the effects of chemogenetic silencing were most visible in network event structure rather than in bulk correlation measures. Given its decorrelating effect within V1, we had expected changes in correlations between V1 and HVAs during periods of high acetylcholine, potentially contributing to the refinement of inter-areal circuits ^49^. The absence of such an effect suggests that inter-areal coupling at this age is still dominated by retinal wave-driven thalamocortical inputs and developing cortico-cortical projections ^5,38^.

What network mechanisms might mediate acetylcholine-driven decorrelation? We confirmed that phasic pyramidal responses to acetylcholine are already heterogeneous at this stage, including a transient hyperpolarization likely mediated by muscarinic receptors as in adult mice ^51,52^. However, the cellular substrates of circuit desynchronization remain to be established, and somatostatin (SST) interneurons are a strong candidate. These interneurons are among the earliest inhibitory populations to mature in the cortex and contribute to the developmental transition from synchronized to desynchronized activity in both V1 and somatosensory cortex ^62,63^. In adult V1, acetylcholine excites SST interneurons through muscarinic M1/M3 receptors, and SST activation is sufficient to decorrelate cortical activity ^64,65^. One plausible mechanism is that basal forebrain acetylcholine recruits SST interneurons via M1/M3 receptors during the second postnatal week, and the resulting dendritic inhibition of pyramidal cells decorrelates ongoing activity. Vasoactive intestinal peptide (VIP) interneurons, which are strongly depolarized by nicotinic and muscarinic signaling in adult V1 ^66,67^, may contribute a complementary disinhibitory drive during movement, although their role at this age remains unknown. A small local contribution from cortical VIP-ChAT interneurons ^68^ cannot be excluded, but these cells are very sparse in the developing visual cortex ^57^. Alternatively or in parallel, muscarinic M2 receptors expressed presynaptically on pyramidal neurons ^69^ could reduce recurrent glutamate release and further weaken lateral excitation ^22^. Our atropine results are consistent with a broad muscarinic contribution to V1 network maturation, in line with previous work using genetic deletion of muscarinic receptor subtypes ^70^. Resolving how these mechanisms operate and whether they act simultaneously or sequentially across the second postnatal week will require cell- and receptor-type-specific manipulations.

A further open question is why the developing visual cortex would need cholinergic desynchronization before eye opening. One possible function is to create the conditions for synapse-specific plasticity. Highly synchronized activity, such as bursts driven by retinal waves, can co-activate neighboring neurons and favor non-selective potentiation. By contrast, brief epochs of decorrelated activity permit the differential coactivity required for Hebbian competition and synaptic refinement ^71,72^. A complementary, non-exclusive role would be homeostatic: periodic interruption of highly synchronous events may prevent overexcitation and support the maturation of excitation-inhibition balance through recruitment of local interneurons ^62,63^. Episodic cholinergic desynchronization may therefore contribute directly to cortical maturation rather than simply reflect the emerging arousal system.

Our study places acetylcholine within the small set of neuromodulators known to shape cortical network activity before eye opening, alongside serotonin ^25^ and oxytocin ^24^. These findings also indicate a distinctive developmental role for acetylcholine that, unlike in adults, links active behavioral states to a transient reduction in V1 activity. How other neuromodulators contribute, and whether disruption of these systems impairs sensory processing or contributes to neurodevelopmental disorders, remain open questions. By coupling behavioral states to the modulation of cortical activity a full week before eye opening, cholinergic signaling may provide an internal instructive signal that helps prepare visual cortex circuits for the transition to sensory-driven processing.

## METHODS

### Animals

We used neonatal C57BL/6J (Janvier) and ChAT-IRES-Cre (JAX #006410) mice between postnatal days 8 and 14 (P8-14), plus one P15 animal included in the P8-14 cohort. Eye opening in C57BL/6J mice occurs on average around P13-14, with some inter-animal variability ^73^. We confirmed that all animals, including the P15 pup, had closed eyelids at the start of the experiment. Therefore, for simplicity, we refer to this cohort as P8-14 throughout. C57BL/6J litters were obtained from timed-pregnant females. ChAT-IRES-Cre litters were bred in-house, and the sire was removed during the first days after birth to improve pup survival. A total of 43 litters contributed to the P8-10 group, the P11-14 group, or both, with no more than three littermates assigned to the same experimental group. We used 19 litters of ChAT-Cre pups (12 for P8-10 and 13 for P11-14) and 24 litters of C57BL/6J pups (9 for P8-10 and 17 for P11-14). Pups were housed with the dam, and both sexes were included in the experiments. Animals were maintained on a 12-hour light/dark cycle with food and water ad libitum. All experimental procedures were approved by the Institutional Animal Care and Use Committee of the Royal Netherlands Academy of Sciences.

### Viral injections

P0-1 mice were cryoanesthetized by wrapping each pup in aluminum foil and placing it on ice for 5 minutes. The pup was then moved to a stereotaxic adaptor with an integrated cooling system to keep the animal’s body temperature between 0 and 4 °C, and the head was secured between rubber head bars. To target the visual cortex, we made a small incision in the skin over V1, and the coordinates for injection were adjusted relative to vascular lambda (1.4 mm lateral, 0.35 mm anterior, 0.1 mm ventral). A glass pipette (GB100F-10, Science Products) loaded with the viral constructs was mounted on a microinjector (Nanoject III, Drummond Instruments) to puncture the skull and inject a small volume (<100 nL) into the superficial cortical layers. The pipette was left in the brain for 3 minutes and then slowly retracted to reduce the risk of reflux. Pups recovered on a heating pad with bedding and were returned to the dam after full recovery of movement and color. We used a second microinjector (Nanoject II, Drummond Instruments) to perform dual injections in V1 (1.4 mm lateral, 0.35 mm anterior, 0.1 mm ventral) and the basal forebrain (2.7 mm anterior, 0.6 mm lateral, 2.6 mm ventral) following the procedure described above. Because most basal forebrain axons project ipsilaterally ^74^, all injections were carried out in the same hemisphere as the cortical viral injections. For most experiments, we injected a 1:1 mix of adeno-associated viruses (AAVs): AAV9-Syn-NES-jRGECO1a-WPRE-SV40 (Addgene #100854) and AAV9-hSyn-GRAB_ACh3.0 (Addgene #121922). In 4 experiments, we used either AAV9-syn-jGCaMP8m-WPRE (Addgene #162375) or AAV9-hSyn-GRAB_ACh4m (plasmid shared by Yulong Li). The mutant version of the GRAB_ACh3.0 sensor, AAV9-hSyn-GRAB_ACh3.0-mut (plasmid shared by Yulong Li), was used in 2 additional experiments. In the basal forebrain, we injected either AAV9-hSyn-DIO-hM4D(Gi)-mCherry (Addgene #44362-AAV9) or AAV9-hSyn-DIO-mCherry (Addgene #50459). AAV1-syn-DIO-EGFP was also injected into the basal forebrain for immunohistochemistry experiments. AAVs were packaged in-house from plasmids as described previously ^75^.

### Craniotomy for in vivo imaging

Surgery for in vivo imaging was adapted from previous protocols ^62^. Animals between P8 and P14 were anesthetized with 3% isoflurane in 0.5 L/min O_2_ and kept on a heating pad at 37 °C throughout the surgery. After confirming anesthesia by the absence of a pedal withdrawal reflex, lidocaine (Xylocaine 50 mg/g) was applied to the scalp for local analgesia, and sterile saline (0.9% NaCl) was injected subcutaneously on the back of the animal to compensate for fluid loss during the procedure. Isoflurane was then reduced to 1%, and a head plate with a circular opening (Ø 4-6 mm) above the right visual cortex (0.5-2.5 mm rostral from lambda, and 1-3 mm lateral from midline) was attached to the skull with bonding agent and dental cement. A small craniotomy was performed at the center of the head plate, above the visual cortex, using a 30G sterile needle, and the exposed cortical surface was cleared with cortex buffer: 125 mM NaCl, 5 mM KCl, 10 mM HEPES, 2 mM MgSO_4_, 2 mM CaCl_2_, and 10 mM glucose (pH 7.4, ∼ 280 mOsm/L). The head plate was secured to the microscope stage, and a borosilicate glass coverslip (#1 thickness) was placed over the head plate opening and sealed with dental cement to reduce motion artifacts during imaging. Sterile saline was administered subcutaneously for rehydration, isoflurane was turned off, and the animal was left undisturbed for 30 minutes before the imaging session began.

Each in vivo imaging session lasted a maximum of 5 hours, during which the unanesthetized pup was head-fixed to the microscope stage with its body on a heating pad at 37 °C. Widefield epifluorescence and/or two-photon imaging were performed in each session.

### Widefield epifluorescence imaging

Widefield images were acquired with a complementary metal-oxide-semiconductor (CMOS) camera (Moment Photometrics) mounted on a Nikon A1R multiphoton microscope with a 4x/0.13 NA air objective (Nikon CFI Plan Fluor) using NIS-Elements software (Nikon). Images were acquired at 1000×1000 pixels after 2x pixel binning, corresponding to a field of view of 2.25 x 2.25 mm. Epifluorescence excitation was provided by an LED system (pE-2; CoolLED). For sequential dual imaging of the green acetylcholine sensor and red calcium indicator, we used 445 nm (excitation filter, Semrock FF01-442/42) and 565 nm (excitation filter, Semrock FF01-542/20) excitation light, respectively, and a dual-bandpass emission filter (Semrock FF01-512/630) to capture both wavelengths on the same CMOS camera. This configuration produced no observable crosstalk between the green and red channels. For single-channel recordings, single-bandpass excitation and emission filters were used with the above LEDs. In a small subset of experiments, only one channel contributed to the analysis because either a single sensor was injected or only one channel was recorded with sufficient quality. LEDs were synchronized with the camera and sequentially triggered using a custom Arduino trigger box controlled via a LabVIEW interface (Tbox, https://github.com/CALohmann/TBox). Each channel was acquired at a rate of 6-8 Hz. Each recording lasted an average of 5 min.

### Two-photon imaging

Two-photon imaging was performed on a Nikon A1R MP microscope with a 16x/0.8 water-immersion objective (Nikon N16XLWD-PF) and a Ti:Sapphire laser (Chameleon II, Coherent) using NIS-Elements software (Nikon). A field of view in V1 was selected based on the topographic organization of the spontaneous activity patterns under widefield imaging. For recordings in L2/3, the acquisition plane was between 150 and 250 μm below the pia. FOV areas of approximately 200 x 200 μm were used for the analysis of baseline activity and DREADD experiments. FOVs for evoked-activity experiments (ACh puff and sensory stimulation) ranged from 200 x 200 μm to 800 x 800 μm. Recordings were acquired at a rate between 7.7 and 15.3 Hz, and each recording lasted an average of 5 min.

### In vivo pharmacology

To block muscarinic receptors in the cortex, atropine (1 mM, Sigma A0257) was dissolved in cortex buffer, applied to the exposed cortex, and covered with 1.5% agarose dissolved in cortex buffer containing 1 mM atropine. Recordings started 15 min after application. For focal application of acetylcholine chloride (1 mM in cortex buffer, Sigma A2661), a borosilicate glass pipette (3-5 MOhm) containing acetylcholine was positioned near the field of view in layer 2/3 of V1 using a micromanipulator. Three to eight pulses of 100-500 ms duration were delivered using a microinjection system (Toohey Spritzer Pressure System) at 10-15 psi, and were synchronized with image acquisition through the Tbox.

### In vivo image processing and analysis

For two-photon recordings, image stacks acquired with the NIS-Elements software were converted to TIFF format and corrected for motion artifacts using the NoRMCorre algorithm ^76^ implemented in CaImAn (https://github.com/flatironinstitute/caiman). Stacks were rotated so that the top border of the image corresponded to the rostral side of the animal, and the right border to the lateral side. Cell segmentation was then performed using Cellpose-SAM ^77^. ROIs touching image borders were discarded, segmentation quality was visually inspected, and single-frame artifacts (z-score > 7 or < −2) were replaced with the preceding frame’s value. ROIs with artifacts exceeding 1% of recording frames were also discarded.

Calcium and acetylcholine ΔF/F_0_ stacks were generated by subtracting the median baseline fluorescence from each frame and dividing the result by F_0_. Mean calcium traces were extracted from the region of interest (ROI) masks using custom Python scripts. Traces were baseline-corrected using a linear fit and low-pass filtered. Calcium transients were then automatically detected based on a minimum prominence of 5 times the standard deviation of the lowest 50% of values in the trace. ROIs with no transients or a mean peak amplitude below 3 SD of the trace were excluded from analysis to remove inactive neurons and ROI artifacts. For acetylcholine signals, the mean signal across the FOV was used to calculate the ΔF/F_0_ trace, baseline-corrected using the locally weighted scatterplot smoothing (LOWESS) algorithm and low-pass filtered. Acetylcholine transients were detected automatically using the SciPy find_peaks function and a prominence threshold of 1 ΔF/F_0_. For recordings with strong initial bleaching (10-30 s), the initial segment was trimmed accordingly. Frequencies and AUC of acetylcholine and calcium transients were computed from the results of SciPy’s find_peaks function, with AUC computed by integrating each detected transient between its onset and offset. For spatial heterogeneity analysis of the acetylcholine sensor, the FOV was divided into 4 x 4 and 8 x 8 grids, and mean ΔF/F_0_ signals were computed for each subregion, after excluding the lowest 30% of pixel values ^41^. Pairwise correlations were calculated between ΔF/F_0_ traces smoothed with a Gaussian kernel (sigma = 2-3 frames) using Pearson correlation.

Network events were identified from two-photon calcium recordings by converting each neuron’s detected calcium peaks to a binary trace and summing the number of active neurons within a 2-s sliding window. Events were defined as epochs in which more than 20% of recorded neurons were co-active. For each event, synchrony metrics were computed exclusively from neurons active during that event: pairwise Pearson correlations were calculated on smoothed ΔF/F₀ traces (Gaussian kernel, σ = 2-3 frames). Temporal jitter was quantified as the standard deviation, across active neurons within the event window, of the time of each neuron’s first detected calcium peak in the ΔF/F₀ traces. Network events were classified as occurring during acetylcholine transients if they began within the transient and overlapped with at least 10% of the acetylcholine transient duration.

For widefield epifluorescence recordings, image stacks acquired with NIS software were converted to TIFF and downsampled by a factor of two. Calcium and acetylcholine ΔF/F_0_ stacks were generated by subtracting and dividing each frame by the median baseline fluorescence (F_0_). To identify visual areas before eye opening, functional correlation maps were generated following previous studies ^38,49^. Briefly, three seed areas within V1, one in RL, and one in PM were selected (10 x 10 pixels each), and their time courses were computed by averaging pixel values within each seed area across all frames. Next, the Pearson correlation between the time course of each seed area and that of every pixel in the FOV was calculated and displayed as a correlation map. Finally, the three V1 correlation maps were combined into a single RGB image, with each channel representing one map.

The final RGB map reflected the topographic organization of the spontaneous activity patterns and adult-like retinotopic structure. Visual area outlines adapted from published maps ^37^ were rotated and scaled to the widefield FOV using a reference pixel at the most rostral point of V1. An additional ROI of 200 x 200 μm was manually selected within V1 using ImageJ. ΔF/F_0_ was extracted from each area, low-pass filtered, and baseline-corrected using a linear fit. Pearson correlation matrices and cross-correlations were calculated using the extracted mean trace from each area. Peaks were automatically detected using SciPy’s find_peaks function with a threshold of 3 SD above the lowest 50% of values of the trace, and a minimum value of 1 ΔF/F_0_.

Acetylcholine transient onset and offset were defined as the time points before and after each peak, respectively, at which the acetylcholine signal crossed below 40% of the peak amplitude. If a second peak occurred before the signal crossed below this threshold, the second transient was merged with the first. For comparisons of the frequency of V1 calcium transients in Fig. 3F, the acetylcholine peak was set as time zero to capture the temporal dynamics of the calcium decrease associated with body movements. For peri-movement and peri-stimulus analysis, calcium and acetylcholine ΔF/F₀ signals were aligned to the onset of movements or stimuli, and peri-event time histograms were generated from all extracted traces. The peak amplitude was defined as the minimum or maximum of the mean ΔF/F₀ trace per animal within the analysis window, with the direction determined by the sign of the majority of time points in that window.

### Behavioral recordings

During imaging sessions, body movements were recorded using a lightweight motion sensor (3-axis femto accelerometer, ∼100 mg) placed on the back of the pup and lightly secured with surgical tape. The signal therefore integrated any body movement captured by the sensor. Face movements were recorded using an infrared camera (Basler acA1300-30gm) at 25 Hz with a near-infrared (NIR) short-pass dichroic filter (850 nm) to filter out artifacts from the two-photon laser. The motion sensor and infrared camera were synchronized with the imaging acquisition signals using the Tbox device. The voltage output of the motion sensor was digitized at 10 kHz with a Digidata 1550 and recorded with Clampex software.

Face videos were acquired in AVI format with PylonRecorder (software by Friedrich Kretschmer), converted to TIFF, and downsampled to 300 x 300 pixels. In adult mice, acetylcholine dynamics closely track whisking, and full facial movements are similarly informative ^19^. Thus, to extract the motion signal from the face videos, we manually defined an ROI of the full face in ImageJ (or, when microscope light-source artifacts were present, a smaller ROI in the whisker pad). A normalized pixel-intensity trace (ΔF/F_0_) was calculated, with F_0_ defined as the median baseline intensity. To match the temporal resolution of the imaging data, the standard deviation of the face and body sensor traces was computed in non-overlapping windows of duration equal to one imaging frame, yielding a downsampled trace in which each data point corresponds to one imaging frame. This reduced high-frequency baseline noise while preserving the movement-related signals, facilitating subsequent peak detection.

From the motion sensor and face camera traces, peaks were automatically detected based on a minimum prominence of 3 x SD of the baseline. Following criteria adapted from previous studies ^19,44^, peaks were then classified as twitches or prolonged movements based on the number of peaks and inter-peak interval. Isolated peaks with an inter-peak interval > 2 s were labeled as twitches, whereas groups of 3 or more peaks with an inter-peak interval < 1 s were labeled as prolonged movements. Given that prolonged body movements were used as a criterion for inferring active wakefulness (see below), prolonged body movements were labeled as wake movements. For analysis of evoked acetylcholine and calcium responses to movements, twitches and prolonged movements not preceded by at least 2 s of stillness were excluded from analysis.

Movements recorded with the motion sensor and face camera were also used to infer the pup’s behavioral state, following criteria adapted from previous studies ^25,44,78^. Prolonged movements recorded by the body sensor, as defined above, were classified as active wakefulness (awake), whereas twitches recorded by the body sensor were considered active sleep (AS). Body twitches within a 3-s interval were considered part of the same behavioral state. In addition, twitches or bouts of facial twitches were also considered AS periods. Behavioral quiescence (BQ) was defined as the remaining time with no movement recorded by the body and face sensors. We did not classify this state as quiet sleep because delta activity emerges around P11 ^30^. Using these criteria, pups spent approximately 90% of the time in AS or BQ, consistent with previous reports using similar recording conditions ^79^.

### DREADD experiments

For in vivo activation of the inhibitory DREADD hM4Di ^80^, we used Compound 21 dihydrochloride (C21, Tocris #6422) dissolved in sterile saline solution and administered at 5-10 mg/kg. After baseline recordings, C21 was injected subcutaneously (1% volume/weight) on the back of hM4Di and mCherry pups under brief anesthesia (30-60 s) with 1.5% isoflurane. Animals recovered within a few seconds after isoflurane was discontinued, and recordings resumed at least 15 minutes after C21 injection. C21 has been shown to have fewer off-target effects than clozapine-N-oxide (CNO), no back-conversion to clozapine, and stable cerebrospinal fluid (CSF) concentrations 60 min after injection ^81^. For patch-clamp experiments, C21 (5-10 μM) was dissolved in ACSF and bath-applied by perfusing the slices with C21-containing ACSF.

### Sensory stimulation

Whisker stimulation was delivered as an air puff through a borosilicate glass pipette (Ø 1 mm) placed 1 to 5 cm away from the left whiskers. A train of short pulses (50-100 ms, 4-8 repetitions, 30 s interval) was programmed in the Tbox software and sent to a microinjection system (Toohey Spritzer Pressure System) set to <10 psi to avoid aversive responses. This pressure was enough to move the whiskers without causing startle responses, vocalizations, or frenzied movements. For light stimulation, a train of short pulses (50-100 ms, 4-8 repetitions, 30-s intervals) programmed in the Tbox software triggered an LED (8000 K) placed 4-6 cm from the left eyelid. Sensory stimulation was synchronized with imaging and behavioral recordings via the Tbox custom Arduino box, and signals were recorded with Clampex. Evoked responses were classified according to whether movement occurred within 2 s of stimulus onset, and trials with movement in the 2 s preceding stimulus onset were excluded from this analysis.

### Immunohistochemistry

For immunohistochemistry of basal forebrain cholinergic neurons and axons, P0-1 mice were injected into the right basal forebrain with AAV1-syn-DIO-EGFP or AAV9-hSyn-DIO-mCherry. Mice between P8 and P14 were then deeply anesthetized with either an intraperitoneal injection of a lethal dose of pentobarbital or an overdose of isoflurane (5% in 0.5 L/min air). When unresponsive, mice were transcardially perfused through the left ventricle with 5-10 mL of pre-warmed phosphate-buffered saline (PBS; Gibco), followed by 5-10 mL of 4% paraformaldehyde (PFA). Brains were then removed and stored in 1% PFA until sectioning.

Fixed brains were rinsed twice with PBS and embedded in 2% agarose (Invitrogen). Coronal sections of 75-100 μm thickness were prepared using a vibratome (Leica VT1000S) and stored in PBS at 4 °C until immunohistochemistry. Brain sections were permeabilized with 0.5% Triton X-100 and 0.01% sodium azide in PBS (PBS-T) for 30 min, then incubated in blocking solution (5% bovine serum albumin in PBS-T) on a rotary shaker at room temperature for 1 h. Brain slices were then incubated overnight at 4 °C with primary antibodies in PBS-T at the following concentrations: goat anti-ChAT (1:300, Millipore AB144P), chicken anti-RFP (1:500, Rockland 600-901-379), and rabbit anti-GFP (1:500, Invitrogen A11122). The following day, sections were washed three times with PBS, and host-specific secondary antibodies were applied in blocking solution (1:500) and incubated for 1-2 h at room temperature. Brain sections were washed three times in PBS and counterstained with DAPI (5 μg/mL). Sections were washed again in PBS and mounted with mounting medium (Vectashield, VectorLabs) on microscope slides (SuperFrost, Thermo Scientific).

Sections were first imaged at 10x with an epifluorescence microscope (Axio Scan Z1) to confirm expression across entire sections. Antibody-positive sections were then imaged using a confocal microscope (Leica TCS SP5 II) with 20× air and 40× oil objectives. Confocal images of the basal forebrain were acquired as single optical planes or as 5 μm z-stacks from 400-700 μm square fields of view. Images of V1 were acquired as 250-300 μm square fields of view in z-stacks spanning 10 μm with a 0.5-1 μm step size. For colocalization analysis, images of single axons were also acquired with a 40× oil objective. Acquisition settings were kept constant across conditions within each experiment.

Images were processed and analyzed in ImageJ (NIH). Maximum-intensity projections of z-stacks were generated, a median filter (1-pixel radius) was applied, and contrast was uniformly adjusted across all images used for quantification. Cholinergic cell counts in the basal forebrain were normalized to area (mm²). In the visual cortex, DAPI labeling and distance from the pial surface were used to delineate layers 1 and 2/3. To isolate varicosity fluorescence from cholinergic axons above background, images were thresholded and quantified using particle analysis in ImageJ. Axon density was defined as the percentage of area covered within layers 1-3 (up to 300 μm from the pia). Three to five ROIs per animal were selected for quantification.

### Slice patch-clamp electrophysiology

Pups were euthanized by decapitation, and brains were quickly removed and immersed in ice-cold oxygenated sucrose cutting solution containing (in mM): 215 sucrose, 2.5 KCl, 1.25 NaH₂PO₄, 26 NaHCO₃, 20 glucose, 1 CaCl₂, and 7 MgCl₂ (pH 7.4, osmolarity adjusted to ≈320 mOsm) bubbled with carbogen gas (95% O₂, 5% CO₂). Coronal slices (300 μm thick) were cut using a vibratome (Leica VT1200S) and transferred to a holding chamber with artificial cerebrospinal fluid (ACSF) at 32 °C containing (in mM): 125 NaCl, 3.5 KCl, 1.25 NaH₂PO₄, 1 MgCl₂, 26 NaHCO₃, 20 glucose, and 2 CaCl₂ (pH 7.4, osmolarity adjusted to 300-310 mOsm) bubbled with carbogen gas (95% O₂, 5% CO₂). Slices were kept at 32 °C for 30 minutes and then transferred to room temperature for at least 30 minutes before use.

Slices were perfused with oxygenated ACSF in a recording chamber at 1 mL/min and maintained at 32 °C. Pyramidal neurons in V1 were identified by their morphology and electrophysiological properties, whereas basal forebrain cholinergic neurons were identified by mCherry fluorescence using an Olympus BX51WI epifluorescence microscope. Whole-cell current-clamp recordings were performed as previously described ^82^ with a MultiClamp 700B amplifier (Molecular Devices), digitized at 20 kHz, and low-pass filtered at 10 kHz using a Digidata 1440 and Clampex 10 software (Molecular Devices). Patch electrodes (3-6 MΩ) were pulled from borosilicate glass (1.5 mm OD, Science Products) and filled with a potassium gluconate (K-gluconate) internal solution containing (in mM): 122 K-gluconate, 13 KCl, 10 HEPES, 10 phosphocreatine-Na, 4 ATP-Mg, and 0.3 GTP-Na, adjusted to pH 7.3 with KOH.

Cells were excluded from analysis if series resistance exceeded 30 MΩ, changed by more than 25% during the recording, or if the membrane potential was more depolarized than -50 mV. Series resistance was compensated, and no correction was applied for the liquid junction potential. Current-clamp recordings were performed without synaptic blockers. Active membrane properties were recorded in current-clamp mode at resting membrane potential and at a holding potential of -60 mV. The number of action potentials was measured in response to depolarizing 500-ms current injections in 10 pA increments. Changes in membrane potential after acetylcholine application (1 mM in ACSF) were measured relative to the mean membrane potential during the 5 s preceding application. Offline analysis was performed by loading the files with pyABF (https://github.com/swharden/pyABF) and analyzing the action potentials with Intrinsic Physiology Feature Extractor (IPFX, https://github.com/AllenInstitute/ipfx).

### Live confocal imaging of acute brain slices

For live confocal imaging, neonatal pups were injected with AAV9-hSyn-GRAB_ACh3.0, and acute brain slices were prepared between P8 and P14 as described above. Slices were placed in the confocal recording chamber and continuously perfused with oxygenated ACSF at 32 °C. Imaging was performed on a Leica SP5 confocal microscope with a 20× water-immersion objective (0.8 NA, Leica). The acetylcholine sensor signal was acquired using an argon laser at 488 nm at a frame rate of 3-5 Hz. Acetylcholine chloride (Sigma A2661), muscarine chloride (Sigma M6532), atropine (Sigma A0257), and glutamate (Sigma G1626) were each dissolved in ACSF and loaded into a borosilicate glass pipette mounted on a micromanipulator and positioned just above the brain slice. One to three pulses of 50-100 ms duration were applied per recording using a microinjection system (Toohey Spritzer Pressure System) at 10-15 psi. For analysis, a circular ROI around the pipette tip was manually defined in ImageJ, and ΔF/F_0_ was calculated, where F_0_ was the median fluorescence of the trace before drug application.

For simultaneous live confocal imaging and patch-clamp electrophysiology, whole-cell current-clamp recordings were performed as described above using a MultiClamp 700B amplifier (Molecular Devices), digitized at 20 kHz, and low-pass filtered at 10 kHz using a Digidata 1440 and Clampex 10 software (Molecular Devices).

### Statistics

Statistical comparisons were performed at the animal level, unless otherwise indicated in the figure legends. Normality was assessed using the Shapiro-Wilk test to determine whether parametric or nonparametric analyses were appropriate. For small sample sizes ≤ 5, parametric tests were applied by default. For two groups, paired data were analyzed using paired t-tests or Wilcoxon signed-rank tests, whereas unpaired data were analyzed using unpaired t-tests or Mann-Whitney U tests. Comparisons among three or more groups were performed using repeated-measures ANOVA or Friedman tests for paired designs, and one-way ANOVA or Kruskal-Wallis tests for unpaired designs. When omnibus tests were significant, multiple comparisons were performed using Tukey, Dunn, or Bonferroni post hoc tests, as appropriate. Statistical analyses were performed in Python using the SciPy and statsmodels libraries; two-way ANOVAs were performed in GraphPad Prism. All data are presented as mean +/- standard error of the mean (SEM); individual points or lines represent individual animals or neurons, as indicated in each figure.

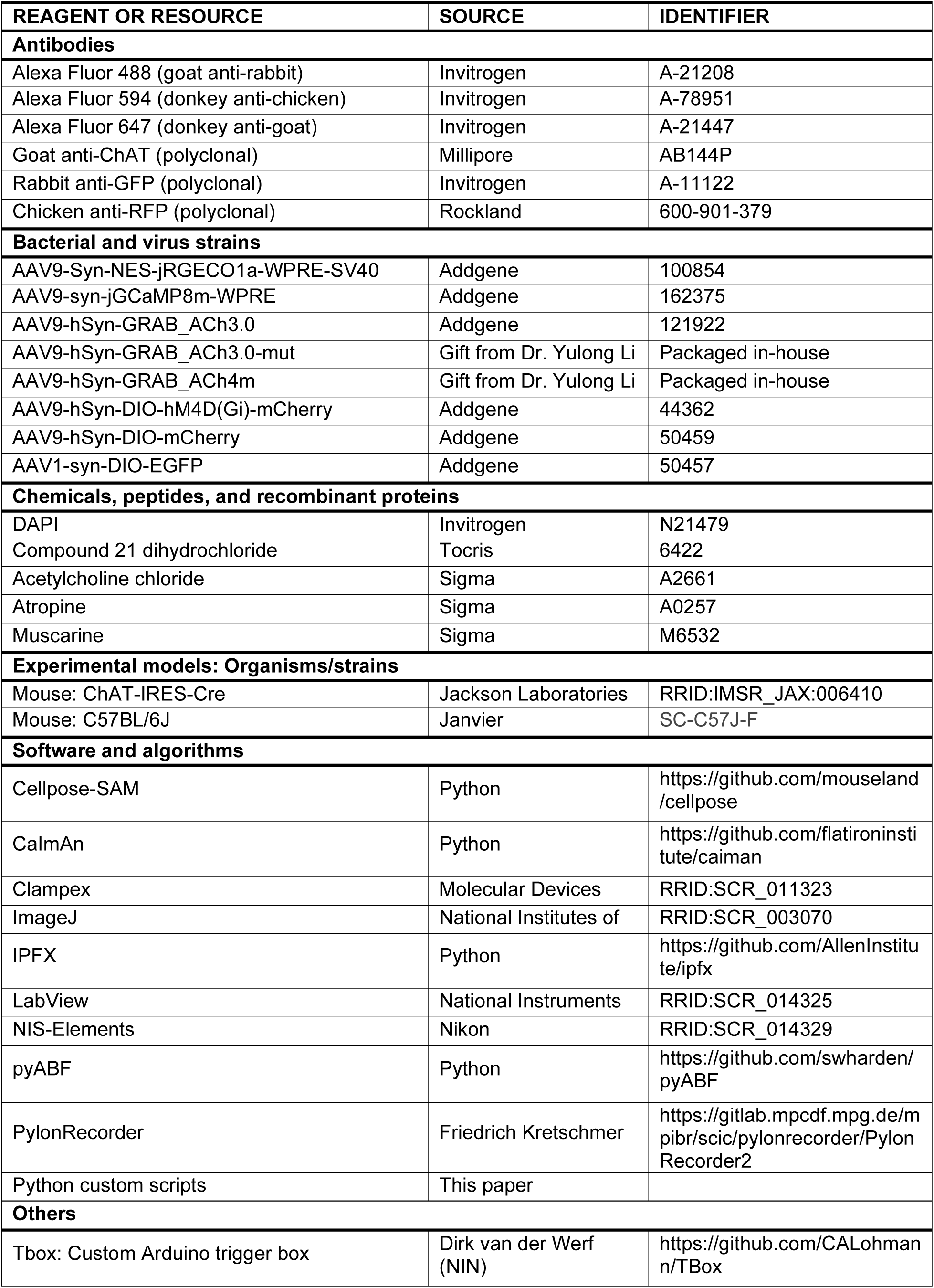

## ACKNOWLEDGMENTS

We thank members of the Lohmann laboratory for helpful discussions during this study, and Renske de Korte for assistance with management of the mouse colony. We also thank Francesca Siclari, Mark Blumberg, and Davide Warm for critically reading the manuscript. We are grateful to Yulong Li (Peking University) for sharing acetylcholine sensor plasmids, to Fred de Winter (NIN) for AAV packaging, and to Friedrich Kretschmer (Max Planck Institute for Brain Research) for adapting the software for video camera recordings. We thank Dirk van der Werf for building the trigger box and the NIN Mechatronics department for technical support with experimental setups. This study was supported by ENW Open Competition grant no. OCENW.KLEIN.535 and Health-Holland grant no. LSHM24002.

## AUTHOR CONTRIBUTIONS

Conceptualization, funding acquisition, supervision, writing-review & editing, C.L.; conceptualization, experiments, analysis, writing-original draft, writing-review & editing, D.C.-G.; experiments, J.A.

---

**Figure S1.**
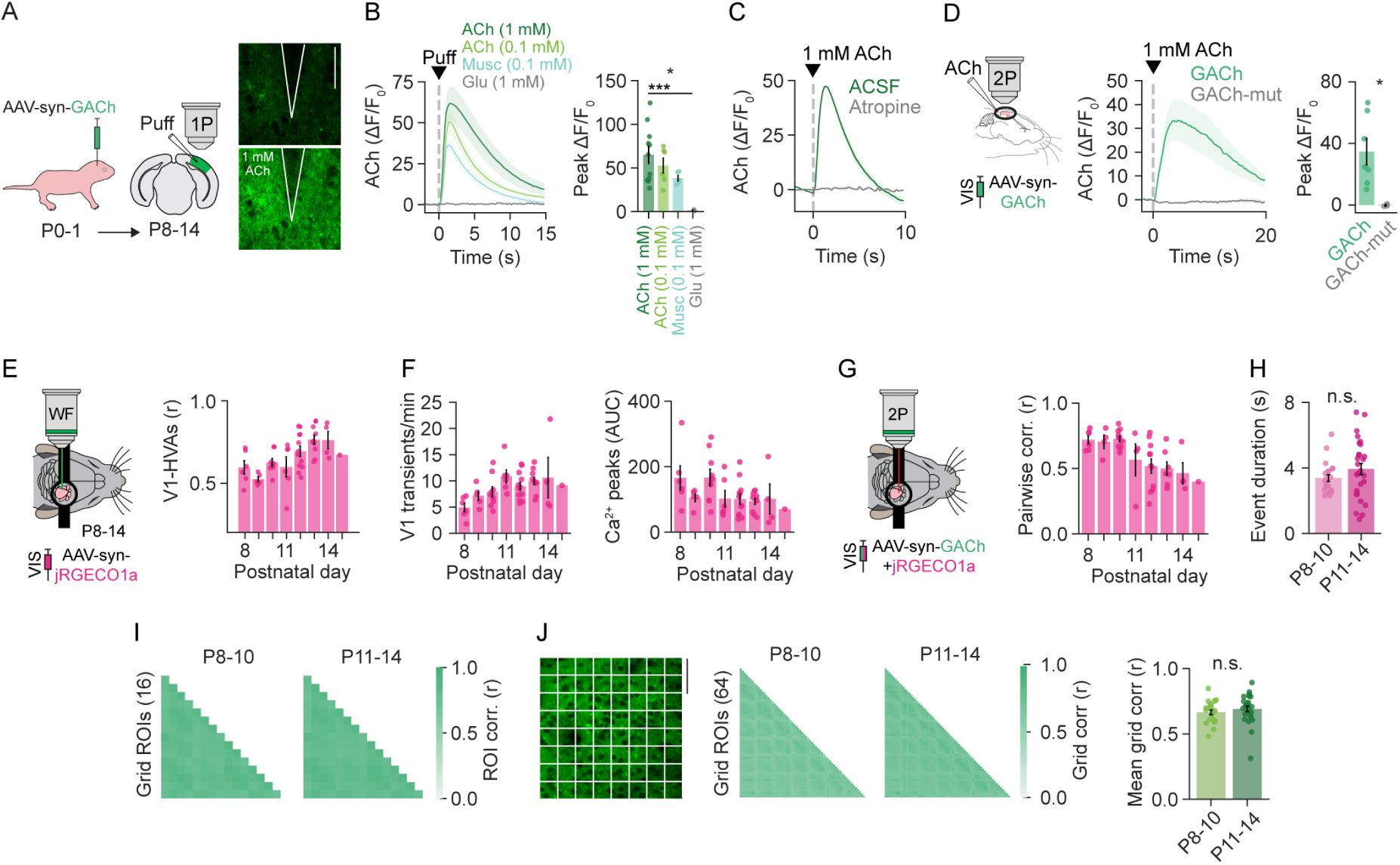
Validation of the acetylcholine sensor and developmental changes in spontaneous network activity during the second postnatal week. **A.** Left: Experimental timeline for AAV-syn-GACh injection at P0-1 and preparation of acute brain slices (P8-14) containing V1 for confocal imaging. Right: Representative frames of a field of view (FOV) in V1 before (top) and after (bottom) application of 1 mM ACh. Scale bar: 50 μm. **B.** Left: GACh time course after application of 1 mM acetylcholine (ACh; n = 11 trials, 6 mice), 0.1 mM ACh (n = 6 trials, 3 mice), muscarine (musc; n = 5 trials, 3 mice), and glutamate (Glu; n = 2 trials, 2 mice). Right: Peak amplitudes of the GACh sensor responses. Whereas application of 1 mM ACh evoked a large fluorescence response, glutamate did not evoke a response. Each dot represents the average response to 1-3 puffs. **C.** Bath application of atropine (10 µM) blocked the response to 1 mM ACh (2 trials, 1 mouse). **D.** In vivo two-photon validation of GACh. 1 mM ACh was puffed onto the superficial layers of V1. Fluorescence increases following ACh puff were compared between the functional acetylcholine sensor (GACh: n = 7 trials, 4 mice) and a nonresponsive mutant version (GACh-mut; n = 3 trials, 1 mouse). **E.** Left: Schematic of in vivo widefield calcium imaging of V1 and HVAs. Right: Correlation of neuronal activity between V1 and HVAs increased during the second postnatal week. **F.** Frequency (left) and AUC (right) of calcium transients in V1 during the second postnatal week. **G.** Left: Schematic of in vivo two-photon calcium imaging of V1. Right: Pairwise neuronal correlations within V1 decreased with age. **H.** Network-event duration did not change between P8-10 (21 mice) and P11-14 (27 mice). **I.** Pearson correlation matrix of a 4 x 4 ROI grid at P8-10 and P11-14, quantified in Fig. 1L. **J.** Left: Example two-photon image of GACh-labeled neurons in layer 2/3 of V1 with an overlaid grid of 64 ROIs. Middle: Pearson correlation matrices among the 64 ROIs at P8-10 and P11-14. Right: Mean grid correlation values per experiment. Data are shown as mean +/- SEM. Each dot represents the mean per animal, unless otherwise stated. For statistical comparisons, the Mann-Whitney test was used in (**H**, **J**), and one-way ANOVA with post hoc Bonferroni was used in (**B**). n.s., not significant; \**P* < 0.05, \*\**P* < 0.01, \*\*\**P* < 0.001, \*\*\*\**P* < 0.0001.

**Figure S2.**
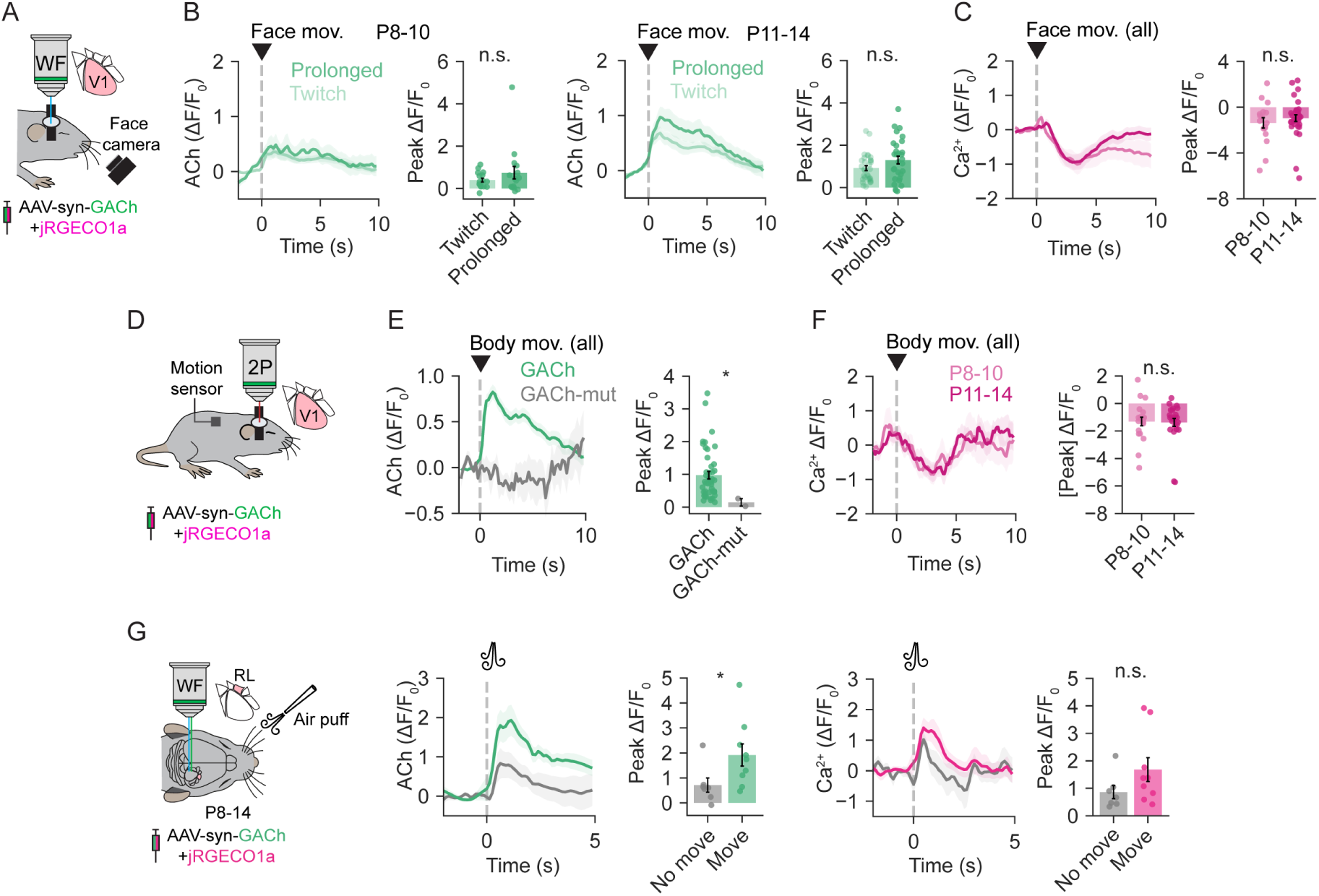
Movement-associated acetylcholine and calcium signals in V1, and whisker-evoked signals in RL. **A.** Twitches and prolonged movements were identified from the traces extracted from the face videos (see Methods) and aligned to the V1-averaged widefield acetylcholine signal. **B.** Peri-event time histogram of mean V1 widefield acetylcholine signal aligned to the onset of face twitches and prolonged movements at P8-10 (left, 16 mice) and P11-14 (right, 31 mice). **C.** Peri-event time histogram of mean V1 widefield calcium signal aligned to the onset of face movements at P8-10 (17 mice) and P11-14 (33 mice). **D.** Body movements were recorded during two-photon imaging of acetylcholine and calcium sensors. **E.** Acetylcholine responses were associated with the onset of body movements (GACh: 45 mice). No response was detected with GACh-mut (2 mice). Data from P8-14 mice. **F.** A small decrease in mean V1 calcium amplitude was detected after the onset of body movements at P8-10 (20 mice) and P11-14 (25 mice). **G.** Whisker stimulation evoked neuronal responses and acetylcholine release in area RL regardless of whether the air puff also elicited movements (no movement: 7 mice; movement: 9 mice). Data are shown as mean +/- SEM. Signals of peri-event and peri-stimulus histograms were normalized to a 2-s pre-event baseline. Peak amplitudes in bar plots were measured within the first 5 s after movement or stimulus onset. Each dot represents the mean for a single animal. Statistical comparisons between two groups were made using unpaired t-tests or Mann-Whitney U tests. n.s., not significant, \**P* < 0.05, \*\**P* < 0.01, \*\*\**P* < 0.001, \*\*\*\**P* < 0.0001.

**Figure S3.**
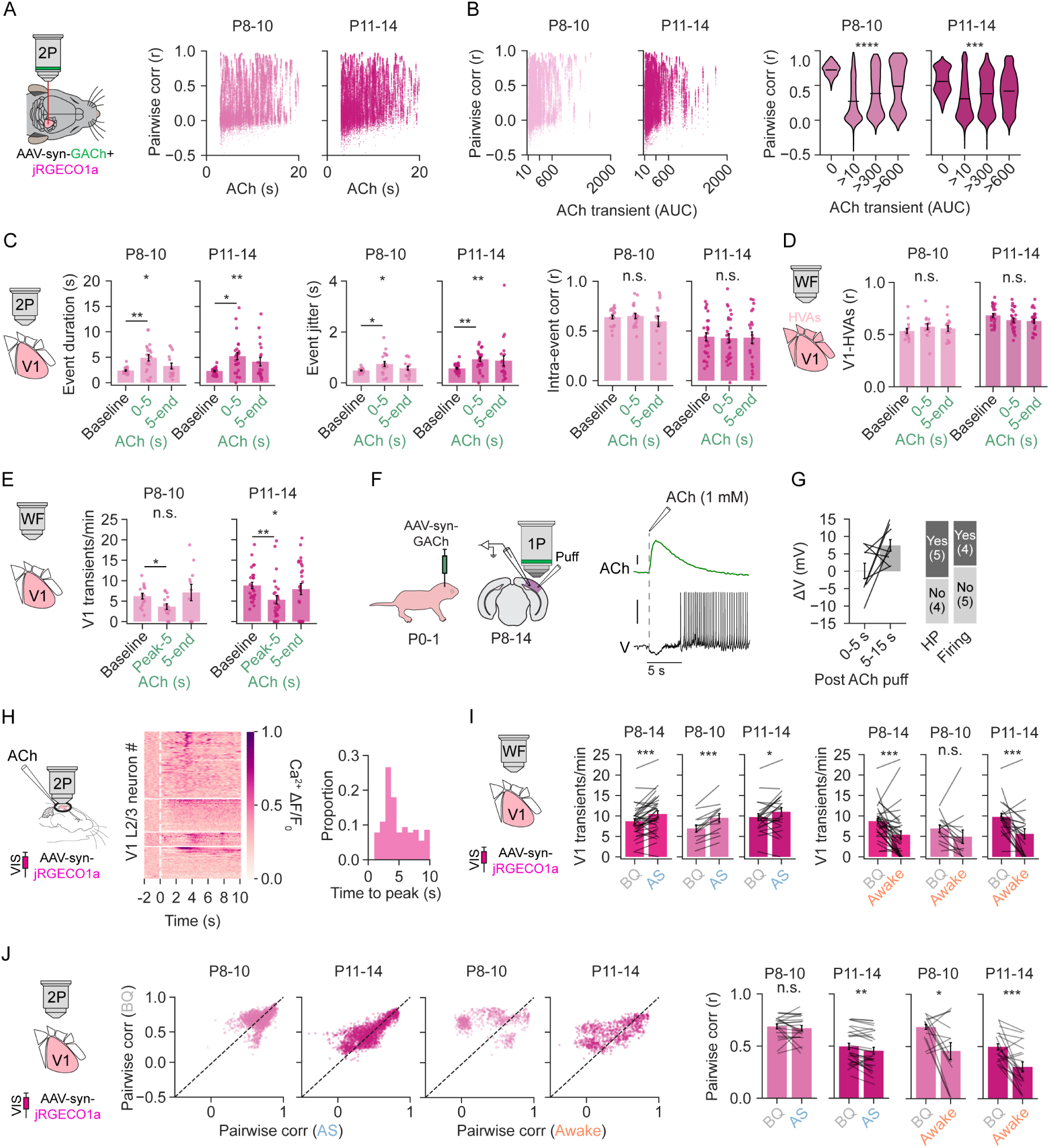
Network and cellular effects of cholinergic signaling and behavioral states on neuronal activity in developing V1. **A.** Relationship between neuronal pairwise correlations and the duration of each acetylcholine transient. Each data point represents one neuron’s mean Pearson correlation with all other neurons during a single acetylcholine transient, plotted against transient duration; neurons from different recordings can therefore align vertically at the same duration. **B.** Left: Relationship between neuronal pairwise correlations and the AUC of each acetylcholine transient. Right: The scatter plot data on the left binned by AUC. P8-10 (baseline: 21 mice; >10: 21 mice; >300: 19 mice; >600: 14 mice); P11-14 (baseline: 27 mice; >10: 27 mice; >300: 26 mice; >600: 21 mice). **C.** Event duration (left), jitter (middle), and intra-event pairwise correlations (right) of synchronized network events during baseline and acetylcholine transients recorded with two-photon microscopy. Data are separated by age group: P8-10 (baseline: 20 mice; 0-5: 20 mice; 5-end: 15 mice) and P11-14 (baseline: 27 mice; 0-5: 26 mice; 5-end: 20 mice). **D.** Correlated activity between V1 and HVAs recorded with widefield imaging was calculated during the first 5 s after onset and during the remainder of the acetylcholine transients. **E.** Frequency of V1 calcium transients during baseline, the first 5 s after the acetylcholine peak, and the rest of the acetylcholine transient by age group. **F.** Left: AAV encoding GACh was injected at P0-1, followed by patch-clamp recording and/or acetylcholine-sensor imaging in acute slices at P8-14. Right: Representative traces of the fluorescence signal of the acetylcholine sensor (top) and a simultaneous whole-cell voltage recording of a pyramidal neuron in response to 1 mM ACh. **G.** Left: Pyramidal neurons in V1 (n = 9, 5 mice) showed heterogeneous voltage and firing responses during the early (0-5 s) and delayed (5-15 s) periods after the acetylcholine puff. Right: Number of neurons showing early hyperpolarization (HP) and delayed firing responses. **H.** In vivo application of 1 mM acetylcholine during two-photon imaging of jRGECO1a-expressing V1 L2/3 neurons (143 neurons, 4 mice). Acetylcholine activated a subset of neurons, with peak response times occurring most frequently at 3-4 s. **I.** Within the same animals, the frequency of V1 calcium transients was compared between BQ and wakefulness and between BQ and AS, both across P8-14 and separately at P8-10 (13 mice in both comparisons) and P11-14 (22 mice in both comparisons). Behavioral states (BQ, AS, awake) were defined from twitches and prolonged movements detected in body and face recordings (see Methods). **J.** Pairwise correlations of neuronal activity in V1 between BQ and AS (left), and BQ and wakefulness (right) at P8-10 (21 mice in both comparisons) and P11-14 (27 mice in both comparisons). Bar plots show mean data per animal from scatter plot data. Pairwise correlations were lower during both AS and wakefulness at P11-14. Data are shown as mean +/- SEM. Unless otherwise noted, each dot represents the mean from a single animal. **D,** P8-10 (Baseline: 15 mice; 0-5: 15 mice; 5-end: 11 mice); P11-14 (Baseline: 26 mice; 0-5: 26 mice; 5-end: 26 mice). **E,** P8-10 (Baseline: 15 mice; Peak-5: 15 mice; 5-end: 11 mice); P11-14 (Baseline: 26 mice; Peak-5: 26 mice; 5-end: 26 mice). One-way ANOVA or Kruskal-Wallis test followed by post hoc Dunn or Bonferroni tests were used for statistical comparisons in **B**-**E**; paired t-tests or Wilcoxon signed-rank tests were used for **I**-**J**. n.s., not significant; \**P* < 0.05, \*\**P* < 0.01, \*\*\**P* < 0.001, \*\*\*\**P* < 0.0001.

**Figure S4.**
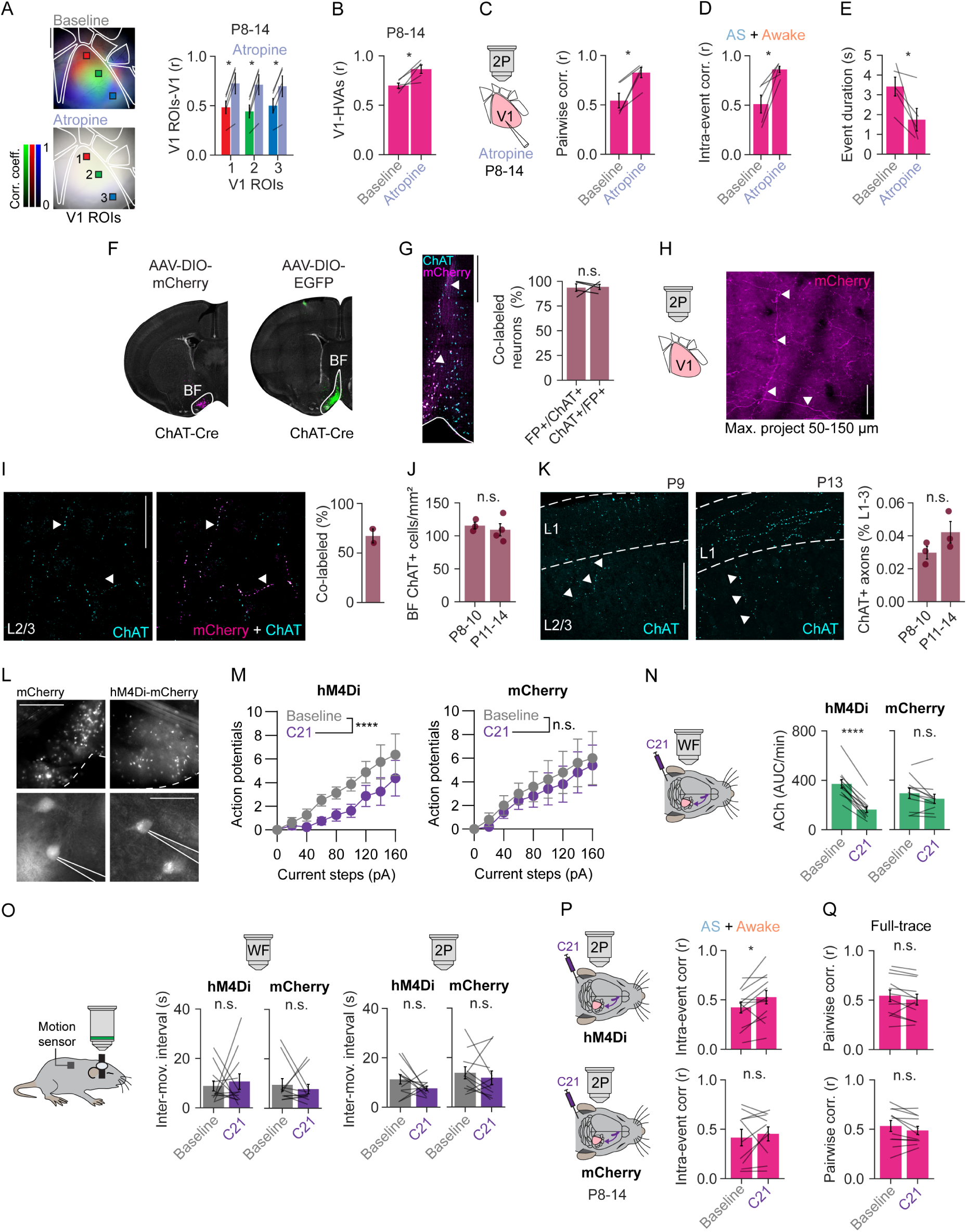
Pharmacological and chemogenetic manipulation of cholinergic signaling in vitro and in vivo. **A.** Local atropine application increased functional correlations between individual V1 ROIs and the rest of V1 at P8-10 and P11-14. Scale bar: 0.5 mm. **B.** Functional correlations between V1 and HVAs increased after atropine application. **C.** Full-trace pairwise correlations in V1 before and after atropine application. **D.** Intra-event pairwise correlations during active behavioral states increased after atropine application. **E.** Network event duration in V1 recorded with two-photon microscopy decreased after atropine application. **F.** Expression of AAV-hSyn-DIO-mCherry (left) and AAV-syn-DIO-EGFP (right) in ChAT-Cre mice was restricted to the basal forebrain (BF), mainly the vertical and horizontal diagonal bands. **G.** Left: Basal forebrain neurons injected with AAV-hSyn-DIO-mCherry and labeled with anti-RFP and anti-ChAT antibodies. White arrowheads point to co-labeled neurons and projecting axons. Scale bar: 500 μm. Right: Colocalization between fluorescent protein (FP) labeling and ChAT expression in basal forebrain neurons (4 mice). **H.** Two-photon image of AAV-DIO-mCherry-labeled cholinergic axons in V1. White arrowheads point to axonal varicosities. Scale bar: 200 μm. **I.** Colocalization between mCherry-labeled axons and ChAT-positive fibers in V1 (2 mice). Scale bar: 50 μm. **J.** Density of basal forebrain cholinergic neurons at P8-10 (3 mice) and P11-14 (4 mice). **K.** Density of ChAT+ axons in L1-3 of V1 at P8-10 (3 mice) and P11-14 (3 mice). White arrowheads point to labeled axonal varicosities. Scale bar: 100 μm. **L.** Brightfield and epifluorescence imaging during whole-cell patch-clamp recordings from hM4Di-mCherry+ and mCherry+ basal forebrain neurons. **M.** Bath application of C21 reduced evoked action potentials in hM4Di (n = 8 neurons, 3 mice) but not in mCherry control neurons (n = 5 neurons, 3 mice) held at -60 mV. **N.** Under widefield imaging, acetylcholine levels (AUC/min) in V1 were reduced after C21 injection in hM4Di mice, but not in mCherry controls. **O.** The inter-event interval of body movements did not change after C21 in hM4Di or mCherry mice. **P.** Intra-event pairwise correlations increased after C21 injection in hM4Di mice during active behavioral states (AS + awake). **Q.** In two-photon recordings, full-trace pairwise correlations were similar after C21 administration in hM4Di and mCherry mice. Data are shown as mean +/- SEM. Unless otherwise noted, each dot represents the mean for a single animal. **A**-**E**: 5 mice, **N-Q**: hM4Di (12 mice), mCherry (11 mice). Paired t-tests were used in **A-E** and **G**, two-way ANOVA in **M**, unpaired t-tests in **J**, **K**, and paired t-tests or Wilcoxon signed-rank tests were used in **N**-**Q**. n.s., not significant; \**P* < 0.05, \*\**P* < 0.01, \*\*\**P* < 0.001, \*\*\*\**P* < 0.0001.

## REFERENCES

1. Huberman, A.D., Feller, M.B., and Chapman, B. (2008). Mechanisms Underlying Development of Visual Maps and Receptive Fields. Annu.Rev Neurosci 31, 479–509.

2. Khazipov, R., and Luhmann, H.J. (2006). Early patterns of electrical activity in the developing cerebral cortex of humans and rodents. Trends in Neurosciences 29, 414–418.

3. Leighton, A.H., and Lohmann, C. (2016). The Wiring of Developing Sensory Circuits-From Patterned Spontaneous Activity to Synaptic Plasticity Mechanisms. Front Neural Circuits 10, 71. 10.3389/fncir.2016.00071.

4. Wu, M.W., Kourdougli, N., and Portera-Cailliau, C. (2024). Network state transitions during cortical development. Nat Rev Neurosci 25. 10.1038/s41583-024-00824-y.

5. Ackman, J.B., Burbridge, T.J., and Crair, M.C. (2012). Retinal waves coordinate patterned activity throughout the developing visual system. Nature 490, 219–225.

6. Burbridge, T.J., Xu, H.-P., Ackman, J.B., Ge, X., Zhang, Y., Ye, M.-J., Zhou, Z.J., Xu, J., Contractor, A., and Crair, M.C. (2014). Visual Circuit Development Requires Patterned Activity Mediated by Retinal Acetylcholine Receptors. Neuron 84, 1049–1064. 10.1016/j.neuron.2014.10.051.

7. Cang, J., Renteria, R.C., Kaneko, M., Liu, X., Copenhagen, D.R., and Stryker, M.P. (2005). Development of Precise Maps in Visual Cortex Requires Patterned Spontaneous Activity in the Retina. Neuron 48, 797–809.

8. Ge, X., Zhang, K., Gribizis, A., Hamodi, A.S., Sabino, A.M., and Crair, M.C. (2021). Retinal waves prime visual motion detection by simulating future optic flow. Science 373. 10.1126/science.abd0830.

9. Siegel, F., Heimel, J.A., Peters, J., and Lohmann, C. (2012). Peripheral and central inputs shape network dynamics in the developing visual cortex in vivo. Current Biology 22, 253–258.

10. Weliky, M., and Katz, L.C. (1997). Disruption of orientation tuning in visual cortex by artificially correlated neuronal activity. Nature 386, 680–685.

11. Gribizis, A., Ge, X., Daigle, T.L., Ackman, J.B., Zeng, H., Lee, D., and Crair, M.C. (2019). Visual Cortex Gains Independence from Peripheral Drive before Eye Opening. Neuron 104, 711–723. 10.1016/j.neuron.2019.08.015.

12. Colonnese, M.T., Kaminska, A., Minlebaev, M., Milh, M., Bloem, B., Lescure, S., Moriette, G., Chiron, C., Ben-Ari, Y., and Khazipov, R. (2010). A Conserved Switch in Sensory Processing Prepares Developing Neocortex for Vision. Neuron 67, 480–498.

13. McCormick, D.A., Nestvogel, D.B., and He, B.J. (2020). Neuromodulation of Brain State and Behavior. Annu. Rev. Neurosci. 43, 391–415. 10.1146/annurev-neuro-100219-105424.

14. Lohani, S., Moberly, A.H., Benisty, H., Landa, B., Jing, M., Li, Y., Higley, M.J., and Cardin, J.A. (2022). Spatiotemporally heterogeneous coordination of cholinergic and neocortical activity. Nat Neurosci 25, 1706–1713. 10.1038/s41593-022-01202-6.

15. Neyhart, E., Zhou, N., Munn, B.R., Law, R.G., Smith, C., Mridha, Z.H., Blanco, F.A., Li, G., Li, Y., Hu, M., et al. (2024). Cortical acetylcholine dynamics are predicted by cholinergic axon activity and behavior state. Cell Reports 43. 10.1016/j.celrep.2024.114808.

16. Reimer, J., McGinley, M.J., Liu, Y., Rodenkirch, C., Wang, Q., McCormick, D.A., and Tolias, A.S. (2016). Pupil fluctuations track rapid changes in adrenergic and cholinergic activity in cortex. Nat Commun 7, 13289. 10.1038/ncomms13289.

17. Jones, B.E. (2020). Arousal and sleep circuits. Neuropsychopharmacol. 45, 6–20. 10.1038/s41386-019-0444-2.

18. Xu, M., Chung, S., Zhang, S., Zhong, P., Ma, C., Chang, W.-C., Weissbourd, B., Sakai, N., Luo, L., Nishino, S., et al. (2015). Basal forebrain circuit for sleep-wake control. Nat Neurosci 18, 1641–1647. 10.1038/nn.4143.

19. Collins, L., Francis, J., Emanuel, B., and McCormick, D.A. (2023). Cholinergic and noradrenergic axonal activity contains a behavioral-state signal that is coordinated across the dorsal cortex. eLife 12, e81826. 10.7554/eLife.81826.

20. Reimer, J., Froudarakis, E., Cadwell, C.R., Yatsenko, D., Denfield, G.H., and Tolias, A.S. (2014). Pupil Fluctuations Track Fast Switching of Cortical States during Quiet Wakefulness. Neuron 84, 355–362. 10.1016/j.neuron.2014.09.033.

21. Yogesh, B., and Keller, G.B. (2024). Cholinergic input to mouse visual cortex signals a movement state and acutely enhances layer 5 responsiveness. eLife 12, RP89986. 10.7554/eLife.89986.5.

22. Goard, M., and Dan, Y. (2009). Basal forebrain activation enhances cortical coding of natural scenes. Nat Neurosci 12, 1444–1449.

23. Pinto, L., Goard, M.J., Estandian, D., Xu, M., Kwan, A.C., Lee, S.-H., Harrison, T.C., Feng, G., and Dan, Y. (2013). Fast modulation of visual perception by basal forebrain cholinergic neurons. Nat. Neurosci. 16, 1857–1863. 10.1038/nn.3552.

24. Maldonado, P.P., Nuno-Perez, A., Kirchner, J.H., Hammock, E., Gjorgjieva, J., and Lohmann, C. (2021). Oxytocin shapes spontaneous activity patterns in the developing visual cortex by activating somatostatin interneurons. Curr Biol 31, 322–333. 10.1016/j.cub.2020.10.028.

25. Ocana-Santero, G., Warming, H., Munday, V., MacKay, H.A., Gibeily, C., Hemingway, C., Stacey, J.A., Saha, A., Lazarte, I.P., Bachetta, A., et al. (2025). Perinatal serotonin signalling dynamically influences the development of cortical GABAergic circuits with consequences for lifelong sensory encoding. Nat Commun 16, 5203. 10.1038/s41467-025-59659-5.

26. Mechawar, N., and Descarries, L. (2001). The cholinergic innervation develops early and rapidly in the rat cerebral cortex: a quantitative immunocytochemical study. Neuroscience 108, 555–567. 10.1016/s0306-4522(01)00389-x.

27. Aubert, I., Cécyre, D., Gauthier, S., and Quirion, R. (1996). Comparative ontogenic profile of cholinergic markers, including nicotinic and muscarinic receptors, in the rat brain. Journal of Comparative Neurology 369, 31–55. 10.1002/(SICI)1096-9861(19960520)369:1%3C31::AID-CNE3%3E3.0.CO;2-L.

28. Peinado, A. (2000). Traveling slow waves of neural activity: a novel form of network activity in developing neocortex. J.Neurosci. 20:RC54, 1–6.

29. Hanganu, I.L., Staiger, J.F., Ben Ari, Y., and Khazipov, R. (2007). Cholinergic Modulation of Spindle Bursts in the Neonatal Rat Visual Cortex In Vivo. J.Neurosci. 27, 5694–5705.

30. Blumberg, M.S., Dooley, J.C., and Tiriac, A. (2022). Sleep, plasticity, and sensory neurodevelopment. Neuron 110, 3230–3242. 10.1016/j.neuron.2022.08.005.

31. Adelsberger, H., Garaschuk, O., and Konnerth, A. (2005). Cortical calcium waves in resting newborn mice. Nat.Neurosci. 8, 988–990.

32. Murata, Y., and Colonnese, M.T. (2018). Thalamus Controls Development and Expression of Arousal States in Visual Cortex. J. Neurosci. 38, 8772–8786. 10.1523/JNEUROSCI.1519-18.2018.

33. Erisken, S., Vaiceliunaite, A., Jurjut, O., Fiorini, M., Katzner, S., and Busse, L. (2014). Effects of Locomotion Extend throughout the Mouse Early Visual System. Current Biology 24, 2899–2907. 10.1016/j.cub.2014.10.045.

34. Niell, C.M., and Stryker, M.P. (2010). Modulation of Visual Responses by Behavioral State in Mouse Visual Cortex. Neuron 65, 472–479.

35. Jing, M., Li, Y., Zeng, J., Huang, P., Skirzewski, M., Kljakic, O., Peng, W., Qian, T., Tan, K., Zou, J., et al. (2020). An optimized acetylcholine sensor for monitoring in vivo cholinergic activity. Nat Methods 17, 1139–1146. 10.1038/s41592-020-0953-2.

36. Dana, H., Mohar, B., Sun, Y., Narayan, S., Gordus, A., Hasseman, J.P., Tsegaye, G., Holt, G.T., Hu, A., Walpita, D., et al. (2016). Sensitive red protein calcium indicators for imaging neural activity. eLife 5, e12727. 10.7554/eLife.12727.

37. Wang, Q., and Burkhalter, A. (2007). Area map of mouse visual cortex. The Journal of Comparative Neurology 502, 339–357. 10.1002/cne.21286.

38. Murakami, T., Matsui, T., Uemura, M., and Ohki, K. (2022). Modular strategy for development of the hierarchical visual network in mice. Nature 608, 578–585. 10.1038/s41586-022-05045-w.

39. Majnik, J., Mantez, M., Zangila, S., Bugeon, S., Guignard, L., Platel, J.-C., and Cossart, R. (2025). Longitudinal tracking of neuronal activity from the same cells in the developing brain using Track2p. Elife 14, RP107540. 10.7554/eLife.107540.

40. Rochefort, N.L., Garaschuk, O., Milos, R.I., Narushima, M., Marandi, N., Pichler, B., Kovalchuk, Y., and Konnerth, A. (2009). Sparsification of neuronal activity in the visual cortex at eye-opening. Proceedings of the National Academy of Sciences 106, 15049–15054.

41. Xuan, F., Li, G., Li, Y., and Dombeck, D.A. (2025). Modulation of speed-dependent acetylcholine release in the hippocampus by spatial task engagement. Cell Reports 44. 10.1016/j.celrep.2025.116443.

42. Disney, A.A., and Higley, M.J. (2020). Diverse Spatiotemporal Scales of Cholinergic Signaling in the Neocortex. J. Neurosci. 40, 720–725. 10.1523/JNEUROSCI.1306-19.2019.

43. Hoy, J.L., and Niell, C.M. (2015). Layer-Specific Refinement of Visual Cortex Function after Eye Opening in the Awake Mouse. J. Neurosci. 35, 3370–3383. 10.1523/JNEUROSCI.3174-14.2015.

44. Del Rio-Bermudez, C., Kim, J., Sokoloff, G., and Blumberg, M.S. (2020). Active Sleep Promotes Coherent Oscillatory Activity in the Cortico-Hippocampal System of Infant Rats. Cerebral Cortex 30, 2070–2082. 10.1093/cercor/bhz223.

45. Chambers, A.R., Kimchi, E.Y., Watanabe, Y., Chakoma, T., and Polley, D.B. (2026). Spontaneous and stimulus-driven arousal produce distinct acetylcholine dynamics across sensory and prefrontal cortex. Preprint at bioRxiv, 10.64898/2026.06.02.729441 https://doi.org/10.64898/2026.06.02.729441.

46. Knudstrup, S.G., Martinez, C., Rauscher, B.C., Doran, P.R., Fomin-Thunemann, N., Kilic, K., Jiang, J., Devor, A., Thunemann, M., and Gavornik, J.P. (2024). Visual stimulation drives retinotopic acetylcholine release in the mouse visual cortex. Preprint at Neuroscience, 10.1101/2024.02.04.578821 https://doi.org/10.1101/2024.02.04.578821.

47. Olcese, U., Iurilli, G., and Medini, P. (2013). Cellular and Synaptic Architecture of Multisensory Integration in the Mouse Neocortex. Neuron 79, 579–593. 10.1016/j.neuron.2013.06.010.

48. Matsumoto, H., Murakami, T., and Ohki, K. (2025). Topographic correspondence between retinotopic and whisker somatosensory map in mouse higher visual area and its development. Front. Neural Circuits 19, 1552130. 10.3389/fncir.2025.1552130.

49. Dwulet, J.M., Zabouri, N., Kirchner, J.H., Wosniack, M.E., Raspanti, A., Kong, D., Houwen, G.J., Maldonado, P.P., Lohmann, C., and Gjorgjieva, J. (2024). Correlated spontaneous activity sets up multi-sensory integration in the developing higher-order cortex. Preprint at bioRxiv, 10.1101/2024.07.19.603239 https://doi.org/10.1101/2024.07.19.603239.

50. Mukherjee, D., Yonk, A.J., Sokoloff, G., and Blumberg, M.S. (2017). Wakefulness suppresses retinal wave-related neural activity in visual cortex. J. Neurophysiol. 118, 1190–1197. 10.1152/jn.00264.2017.

51. Gulledge, A.T., and Stuart, G.J. (2005). Cholinergic Inhibition of Neocortical Pyramidal Neurons. J.Neurosci. 25, 10308–10320.

52. Hedrick, T., and Waters, J. (2015). Acetylcholine excites neocortical pyramidal neurons via nicotinic receptors. Journal of Neurophysiology 113, 2195–2209. 10.1152/jn.00716.2014.

53. Huppé-Gourgues, F., Jegouic, K., and Vaucher, E. (2018). Topographic Organization of Cholinergic Innervation From the Basal Forebrain to the Visual Cortex in the Rat. Frontiers in Neural Circuits 12.

54. Li, X., Yu, B., Sun, Q., Zhang, Y., Ren, M., Zhang, X., Li, A., Yuan, J., Madisen, L., Luo, Q., et al. (2018). Generation of a whole-brain atlas for the cholinergic system and mesoscopic projectome analysis of basal forebrain cholinergic neurons. PNAS 115, 415–420. 10.1073/pnas.1703601115.

55. Thompson, K.J., Khajehali, E., Bradley, S.J., Navarrete, J.S., Huang, X.P., Slocum, S., Jin, J., Liu, J., Xiong, Y., Olsen, R.H.J., et al. (2018). DREADD Agonist 21 Is an Effective Agonist for Muscarinic-Based DREADDs *in Vitro* and *in Vivo*. ACS Pharmacol. Transl. Sci. 1, 61–72. 10.1021/acsptsci.8b00012.

56. Kilb, W., and Luhmann, H.J. (2003). Carbachol-induced Network Oscillations in the Intact Cerebral Cortex of the Newborn Rat. Cereb. Cortex 13, 409–421. 10.1093/cercor/13.4.409.

57. Goral, R.O., Lamb, P.W., and Yakel, J.L. (2024). Acetylcholine neurons become cholinergic during three time windows in the developing mouse brain. eNeuro 11, ENEURO.0542-23.2024. 10.1523/ENEURO.0542-23.2024.

58. Harrison, T.C., Pinto, L., Brock, J.R., and Dan, Y. (2016). Calcium Imaging of Basal Forebrain Activity during Innate and Learned Behaviors. Front. Neural Circuits, 36. 10.3389/fncir.2016.00036.

59. Dominguez, S., Ma, L., Yu, H., Pouchelon, G., Mayer, C., Spyropoulos, G.D., Cea, C., Buzsáki, G., Fishell, G., Khodagholy, D., et al. (2021). A transient postnatal quiescent period precedes emergence of mature cortical dynamics. Elife 10, e69011. 10.7554/eLife.69011.

60. Dooley, J.C., and Blumberg, M.S. (2018). Developmental “awakening” of primary motor cortex to the sensory consequences of movement. eLife 7, e41841. 10.7554/eLife.41841.

61. Laszlovszky, T., Schlingloff, D., Hegedüs, P., Freund, T.F., Gulyás, A., Kepecs, A., and Hangya, B. (2020). Distinct synchronization, cortical coupling and behavioral function of two basal forebrain cholinergic neuron types. Nat Neurosci 23, 992–1003. 10.1038/s41593-020-0648-0.

62. Leighton, A.H., Cheyne, J.E., Houwen, G.J., Maldonado, P.P., De Winter, F., Levelt, C.N., and Lohmann, C. (2021). Somatostatin interneurons restrict cell recruitment to retinally driven spontaneous activity in the developing cortex. Cell Rep 36, 109316. 10.1016/j.celrep.2021.109316.

63. Mòdol, L., Moissidis, M., Selten, M., Oozeer, F., and Marín, O. (2024). Somatostatin interneurons control the timing of developmental desynchronization in cortical networks. Neuron 112, 2015–2030.e5. 10.1016/j.neuron.2024.03.014.

64. Chen, N., Sugihara, H., and Sur, M. (2015). An acetylcholine-activated microcircuit drives temporal dynamics of cortical activity. Nature Neuroscience 18, 892–902. 10.1038/nn.4002.

65. Muñoz, W., Tremblay, R., Levenstein, D., and Rudy, B. (2017). Layer-specific modulation of neocortical dendritic inhibition during active wakefulness. Science 355, 954–959. 10.1126/science.aag2599.

66. Alitto, H.J., and Dan, Y. (2012). Cell-type-specific modulation of neocortical activity by basal forebrain input. Front Syst Neurosci 6, 79. 10.3389/fnsys.2012.00079.

67. Fu, Y., Tucciarone, J.M., Espinosa, J.S., Sheng, N., Darcy, D.P., Nicoll, R.A., Huang, Z.J., and Stryker, M.P. (2014). A Cortical Circuit for Gain Control by Behavioral State. Cell 156, 1139– 1152. 10.1016/j.cell.2014.01.050.

68. Granger, A.J., Wang, W., Robertson, K., El-Rifai, M., Zanello, A.F., Bistrong, K., Saunders, A., Chow, B.W., Nuñez, V., Turrero García, M., et al. (2020). Cortical ChAT+ neurons co-transmit acetylcholine and GABA in a target- and brain-region-specific manner. eLife 9, e57749. 10.7554/eLife.57749.

69. Groleau, M., Kang, J.I., Huppé-Gourgues, F., and Vaucher, E. (2015). Distribution and effects of the muscarinic receptor subtypes in the primary visual cortex. Front Synaptic Neurosci 7. 10.3389/fnsyn.2015.00010.

70. Groleau, M., Nguyen, H.N., Vanni, M.P., Huppé-Gourgues, F., Casanova, C., and Vaucher, E. (2014). Impaired functional organization in the visual cortex of muscarinic receptor knock-out mice. NeuroImage 98, 233–242. 10.1016/j.neuroimage.2014.05.016.

71. Katz, L.C., and Shatz, C.J. (1996). Synaptic activity and the construction of cortical circuits. Science 274, 1133–1138.

72. Winnubst, J., Cheyne, J.E., Niculescu, D., and Lohmann, C. (2015). Spontaneous activity drives local synaptic plasticity in vivo. Neuron 87, 399–410. 10.1016/j.neuron.2015.06.029.

73. Cancedda, L., Putignano, E., Sale, A., Viegi, A., Berardi, N., and Maffei, L. (2004). Acceleration of Visual System Development by Environmental Enrichment. J. Neurosci. 24, 4840–4848. 10.1523/JNEUROSCI.0845-04.2004.

74. Do, J.P., Xu, M., Lee, S.-H., Chang, W.-C., Zhang, S., Chung, S., Yung, T.J., Fan, J.L., Miyamichi, K., Luo, L., et al. (2016). Cell type-specific long-range connections of basal forebrain circuit. eLife 5, e13214. 10.7554/eLife.13214.

75. Pietersz, K.L., Nijhuis, P.J.H., Klunder, M.H.M., Van Den Herik, J., Hobo, B., De Winter, F., and Verhaagen, J. (2024). Production of High-Yield Adeno Associated Vector Batches Using HEK293 Suspension Cells. JoVE, 66532. 10.3791/66532.

76. Pnevmatikakis, E.A., and Giovannucci, A. (2017). NoRMCorre: An online algorithm for piecewise rigid motion correction of calcium imaging data. Journal of Neuroscience Methods 291, 83–94. 10.1016/j.jneumeth.2017.07.031.

77. Pachitariu, M., Rariden, M., and Stringer, C. (2025). Cellpose-SAM: superhuman generalization for cellular segmentation. Preprint at Bioinformatics, 10.1101/2025.04.28.651001 https://doi.org/10.1101/2025.04.28.651001.

78. Gómez, L.J., Dooley, J.C., Sokoloff, G., and Blumberg, M.S. (2021). Parallel and Serial Sensory Processing in Developing Primary Somatosensory and Motor Cortex. J. Neurosci. 41, 3418– 3431. 10.1523/JNEUROSCI.2614-20.2021.

79. Rensing, N., Moy, B., Friedman, J.L., Galindo, R., and Wong, M. (2018). Longitudinal analysis of developmental changes in electroencephalography patterns and sleep-wake states of the neonatal mouse. PLOS ONE 13, e0207031. 10.1371/journal.pone.0207031.

80. Krashes, M.J., Koda, S., Ye, C., Rogan, S.C., Adams, A.C., Cusher, D.S., Maratos-Flier, E., Roth, B.L., and Lowell, B.B. (2011). Rapid, reversible activation of AgRP neurons drives feeding behavior in mice. J. Clin. Invest. 121, 1424–1428. 10.1172/JCI46229.

81. Jendryka, M., Palchaudhuri, M., Ursu, D., van der Veen, B., Liss, B., Kätzel, D., Nissen, W., and Pekcec, A. (2019). Pharmacokinetic and pharmacodynamic actions of clozapine-N-oxide, clozapine, and compound 21 in DREADD-based chemogenetics in mice. Sci Rep 9. 10.1038/s41598-019-41088-2.

82. Li, M., Cabrera-Garcia, D., Harrison, N.L., and Yang, G. (2026). Activation by alcohol of prefrontal layer 5 pyramidal neurons depends on ascending dopaminergic input. J Neurosci, e2105252026. 10.1523/JNEUROSCI.2105-25.2026.

